# Isoform-level resolution in single-cell CRISPR screens reveals hidden functional consequences of gene perturbation

**DOI:** 10.64898/2026.07.09.737410

**Authors:** Nathanael Andrews, Josie Gleeson, Jasper Panten, Sofia Öling, Sofia Lundqvist, Tuuli Lappalainen

**Author notes:** Correspondence (N.A), (J.G), (T.L). Nathanael Andrews and Josie Gleeson contributed equally to this work.

## Abstract

Single-cell CRISPR screens have enabled systematic investigation of gene function, but studies have largely focused on gene-level effects, overlooking transcriptional complexity and isoform usage. Methods capable of capturing splicing and isoform usage have emerged, including long-read sequencing and alternative library preparation strategies, but their suitability for large-scale perturbation screens remains unevaluated. We compare two library preparation methods (10x Genomics and Parse Biosciences) across Illumina short-read, Oxford Nanopore, and PacBio long-read sequencing, applying CRISPRi to silence three genes with distinct regulatory roles (*DDX6, GEMIN5, GFI1B*) in K562 cells. While short-read methods detected some splicing events, only long-read sequencing consistently captured isoform-level changes. Although Parse provided even transcript coverage, we observed strong intronic read enrichment, limiting its utility for splicing analysis. The primary constraint of long-read approaches was sequencing depth: ∼21 million reads are needed for 80% saturation of splicing events in a single perturbation. Notably, *GEMIN5* knockdown produced only modest differential expression but the most extensive splicing changes, an effect invisible to gene-level analysis, underscoring the value of isoform-level screens. We provide a practical framework for isoform-level analysis in single-cell CRISPR screens, identifying current capabilities and limitations. As perturbation studies scale, long-read sequencing will be essential for comprehensive functional interpretation, capturing biology missed by gene-level analysis.

**Graphical Abstract:** 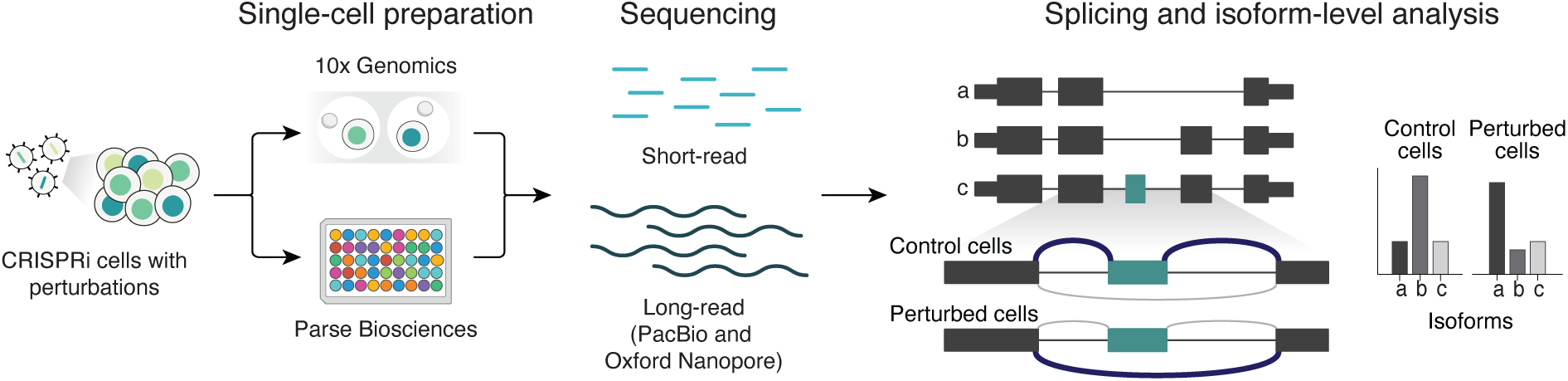

## Background

Single-cell RNA sequencing (scRNA-seq) technologies have become integral tools in functional genomics research. These scRNA-seq methods have been combined with pooled CRISPR screens to perturb genes of interest and assess resulting transcriptional changes at scale, revolutionizing our ability to systematically investigate gene function (1–4). However, single-cell CRISPR screens have predominantly focused on gene expression changes that result from perturbations, overlooking other forms of gene regulation such as alternative splicing and isoform usage (2, 4). Furthermore, the choice of single-cell platform and sequencing technology are key variables in CRISPR screen design, together determining which perturbation effects can be detected and whether the analysis captures transcriptome complexity beyond gene expression (5–7).

Current scRNA-seq platforms differ substantially in their approaches and capabilities. Droplet-based methods, such as 10x Genomics, dominate the field due to their high throughput and established workflows (8). These methods capture either the 5’ or 3’ end of transcripts using oligo-dT priming, resulting in reads concentrated at transcript ends. In contrast, combinatorial barcoding approaches, such as Parse Biosciences’ Evercode, aim to minimize cost and maximize scalability (9). Parse employs random hexamer priming alongside oligo-dT, enabling more even coverages along transcripts and the capture of non-polyadenylated transcripts. However, the split-pool approach requires cell permeabilization during barcoding, which influences which transcripts are ultimately captured. These differences in priming strategy, transcript coverage, and cell handling determine not only transcript representation but also how comprehensively splicing and isoform usage can be assessed (10, 11).

Long-read technologies have emerged as powerful tools for transcriptome analysis. Oxford Nanopore Technologies (ONT) sequences by measuring changes in electrical current as RNA or DNA passes through protein nanopores, translating voltage disruptions into base calls (12). This approach offers flexibility and portability, though with higher error rates compared to short-read methods (13). PacBio’s circular consensus sequencing achieves higher accuracy but with less flexible instrumentation (14). Both platforms have enabled advances in characterizing repetitive regions, reference-free assembly, and resolution of transcript isoforms and splicing patterns, though at substantially lower throughput than short-read sequencing (15–19). As a result, while efforts to integrate long-read sequencing with single-cell CRISPR screens have begun to emerge (20), such studies remain limited in scale and systematic evaluation of platform performance for perturbation screens is lacking (21).

Here, we benchmark common library preparation and sequencing approaches for their ability to capture splicing and isoform changes following single-cell CRISPR perturbation. We find that long-read sequencing reveals perturbation-associated splicing changes undetected by short-read methods, demonstrating the value of isoform-level resolution for comprehensive CRISPR screen analysis.

## Materials and Methods

### CRISPRi K562 cell line construction

The CRISPRi cell line was previously generated by transducing K562 cells (ATCC, CCL243) with lentiCRISPRi(v2)-Blast (Addgene plasmid #170068; http://n2t.net/addgene:170068; RRID:Addgene_170068), which expresses dCas9 fused to KRAB and MeCP2 domains and a blasticidin resistance gene (3). Cells were cultured in K562 culture medium consisting of IMDM (Gibco) with 10% fetal calf serum (Gibco), 1% penicillin and streptomycin (Gibco) and 5 μg / ml blasticidin (Gibco).

### Target gene sgRNA library design and cloning

For each target gene (*GFI1B*, *GEMIN5*, *DDX6*), two guide RNA (gRNA) sequences that showed knockdown efficiencies greater than 50% were taken from Replogle et al (4). For *GFI1B*, a third gRNA was designed following the approach in Domingo et al (49). Finally, five non-targeting control (NTC) gRNAs were chosen, previously used in Replogle et al (4) (**Supplementary Table 6**). The gRNA pool was purchased as an equimolar single-stranded oligo pool (IDT) with 5’/3’ overhangs (5’-TTGTGGAAAGGACGAAACACCG-3’ / 5’-GTTTAAGAGCTATGCTGG-3’) and cloned into the target vector (CROP-seq-opti, Addgene 106280). 10 ng of the pool was amplified using Phusion High-Fidelity PCR Master Mix (BioNordika) with 5’-GGAAAGGACGAAACACCG-3’ and 5’-GCTGTTTCCAGCATAGCTCTTAAAC-3’ (500 nM) (1: 98 °C 30s, 2: 98 °C 10s, 60 °C 30s, 72 °C 30s (2-4 35 cycles), 72 °C 300s, 4 °C) and purified using the QIAGEN PCR purification kit (Qiagen). The target vector was digested using FastDigest Esp3l (Thermo Fisher), and the plasmid backbone was purified using the QIAquick Gel Extraction Kit (QIAGEN). 100ng of the plasmid backbone was assembled with the amplified guide library at a 30x molar excess using 5 reactions of the In-Fusion® Snap Assembly Master Mix (Takara). The assembled product was purified using ethanol precipitation, and 2 μL of this was heat-shock transformed into 5 × 50 μL of NEB Stable Competent E. coli (NEB) according to the manufacturer’s protocol, yielding >300 transformation events per gRNA, followed by plasmid library purification. Representation of gRNAs in the library was confirmed using ONT Lite Whole Plasmid Sequencing (Eurofins Genomics). Virus libraries were produced by transfecting one T75 flask of HEK293FT cells at 80% confluency with a total of 20 μg of the transfer plasmid with equimolar concentrations of pMD2.G (Addgene 12259) and psPAX2 (Addgene 12260) using Lipofectamine 3000 (Invitrogen) in Opti-MEM (Gibco). After 24h, Opti-MEM was replaced with 12ml HEK293FT medium (DMEM, high glucose (Gibco) supplemented with 10% fetal calf serum (Gibco), 2 mM L-glutamine (Gibco), 0.1 mM MEM non-essential amino acids (Gibco), 1 mM Sodium Pyruvate (Gibco) and 500 μg/ml geneticin (Gibco)). Viral supernatants were harvested at 48h and 72h and filtered through a 0.40 μm filter. The filtered supernatants were mixed with 1/3 volume of Lenti-X^TM^ concentrator (Takara), incubated at 4 °C overnight and centrifuged for 60 minutes at 3000g. The resulting lentiviral concentrates were frozen at −20 °C and titrated by transducing serial dilutions of 20 μL into 200, 000 K562 cells and assessing relative survival after puromycin selection (2 μg/ml). The K562-CRISPRi cells were transduced at a multiplicity of infection (MOI) of 0.1, and puromycin selection was initiated at day 1. On day 8, two rounds of dead-cell depletion were performed using the Dead Cell Removal Kit (Miltenyi) according to the manufacturer’s specifications. The resulting cell suspension was directly processed using the 10x Genomics 5’ library kit (10x Genomics) or fixed using the Evercode^TM^ Cell Fixation Kit (Parse Biosciences) and stored at −80 °C.

### 10x Genomics library preparation and sequencing

Cells were processed using the 10x Genomics 5’ library kit (10x Genomics) according to the manufacturer’s instructions with the following modifications: to improve the amplification of full-length transcripts, the reverse transcription (GEM-RT incubation) was increased from 45 minutes to 2 hours, and the cDNA amplification was run with an extension time of 3 minutes rather than 1 minute. For short-read sequencing, pooled libraries were sequenced on NovaSeq X Plus (NovaSeq X Series Control Software 1.3.1.59007) with a 28nt(Read1)-10nt(Index1)-10nt(Index2)-90nt(Read2) setup using a ‘10B’ mode flowcell. The BCL to FASTQ conversion was performed using bcl2fastq_v2.20.0.422 from the CASAVA software suite (Illumina).

For Oxford Nanopore Technologies long-read sequencing, we used the FLT-Seq (v2) protocol as described previously (15), followed by the Ligation Sequencing Kit V14 (SQK-LSK114). The library was sequenced on two PromethION flow cells (FLO-PRO114M). Raw reads were basecalled using dorado with the ‘dna_r10.4.1_e8.2_400bps_sup@v5.0.0’ model.

For PacBio long-read sequencing, we followed the PacBio Kinnex single-cell protocol ‘Preparing Kinnex libraries using the Kinnex single-cell RNA kit’ with no changes using 69ng of cDNA input. The cDNA library was sequenced on a single SMRT cell using the Revio system. The HiFi reads were subsequently segmented using the ‘skera’ tool from PacBio.

### Parse Biosciences library preparation and sequencing

Cells were fixed and permeabilized using the Parse Evercode Cell Fixation Kit, then thawed and loaded into a Parse Evercode Whole Transcriptome (WT) kit v3 (Parse Biosciences), targeting an output of 2000 cells. Cells were processed according to the manufacturer’s specifications and split into two sub-libraries. Libraries of amplified transcripts containing gRNA sequences were generated using the Parse CRISPR Detect kit (Parse Biosciences) according to the manufacturer’s specifications. The resulting WT and CRISPR sublibraries were pooled at a molar ratio of 1:10 and sequenced on a NovaSeq X Plus (NovaSeq X Series Control Software 1.3.1.59007) with a 151nt(Read1)-10nt(Index1)-10nt(Index2)-151nt(Read2) setup using a ‘10B’ mode flowcell. The BCL to FASTQ conversion was performed using bcl2fastq_v2.20.0.422 from the CASAVA software suite (Illumina).

### Data processing and bioinformatic analysis of 10x Genomics Illumina short-read data

The FASTQs were aligned using Cell Ranger ‘count’ (v9.0.1) with the GRCh38.p14 genome and Gencode v48 annotations (8, 50). The gRNA sequences were provided using the feature flag within Cell Ranger to perform guide calling. The pipeline outputs BAM files and count matrices.

### Data processing and bioinformatic analysis of Oxford Nanopore Technologies data

FASTQ files were processed using the nf-core/scnanoseq pipeline (v1.2.1) (51). The FASTQ files for each flow cell were processed separately before being combined to enable a fair comparison with the single SMRT cell sequenced with PacBio. The cell barcodes identified by Cell Ranger with a single guide RNA were used as the custom barcode whitelist. The human reference genome GRCh38.p14 and Gencode v48 annotations were provided for alignment and quantification (50). The scnanoseq pipeline outputs BAM files at each step along with Cell Ranger-style gene and isoform count matrices.

### Data processing and bioinformatic analysis of PacBio data

PacBio segmented reads were processed with the Iso-Seq single-cell workflow (v4.3.0). The cell barcodes identified by Cell Ranger with a single guide RNA were provided as the custom barcode whitelist. Segmented reads were produced using skera (v1.4.0), before cDNA primer and polyA tail trimming. Cell barcodes were extracted and corrected against the custom whitelist. Unique molecular identifiers (UMIs) were clustered by cell barcode and deduplicated using the Iso-Seq ‘groupdedup’ command. The resulting BAM file was processed with the pigeon workflow for transcript mapping, collapsing and classification (v1.4.0). Reads were aligned and collapsed into transcript isoforms using the human reference genome GRCh38.p14 with Gencode v48 annotations (50). Lastly, gene and isoform expression matrices were generated using the pigeon ‘make-seurat’ command.

### Data processing and bioinformatic analysis of Parse Biosciences Illumina short-read data

FASTQs were processed using Parse Biosciences’ split-pipe pipeline (v1.6.4). The two transcriptome cDNA sublibraries were processed separately, followed by their respective CRISPR cDNA sublibraries. The sublibraries were subsequently combined using the pipeline’s ‘--mode comb’ command. The split-pipe script identified and corrected cell barcodes, aligned reads to the human genome GRCh38.p14, assigned transcript counts to genes using Gencode v48 annotations, and generated BAM files and a gene count matrix.

### Gene detection profiles across datasets

RSeQC was used to assess gene coverage with the ‘geneBody_coverage.py’ script and the RSeQC-provided BED file of housekeeping genes, along with one BAM file per dataset (52). Read biotype classification and splice junction quantification were performed using a custom Python pipeline. Gene annotation was parsed from Gencode v48 to extract exon and intron coordinates per gene, classifying genes into categories including protein-coding, lncRNA, pseudogene, mitochondrial, and rRNA. Reads from each dataset’s genome BAM file were assigned to genes based on maximum overlap, with biotype hierarchy as a tiebreaker. Protein-coding and lncRNA reads mapping to multi-exon genes were further classified by splice status — splice junction-spanning, exonic only, or intronic — based on CIGAR string parsing and overlap with annotated intron coordinates. The number of splice junctions per read was recorded for protein-coding multi-exon reads.

### Nuclear-cytosolic enrichment analysis

To examine the enrichment of nuclear or cytosolic genes in the SR datasets, we downloaded public cell fractionation data for the nuclear (doi:10.17989/ENCSR530NHO) and cytosol (doi:10.17989/ENCSR384ZXD) compartments of K562 cells from ENCODE (53). A log_2_ ratio was calculated of the mean (normalized and log transformed) gene expression in the nuclear fraction divided by the cystosolic fraction. Genes with a positive value were more highly expressed in the nuclear compartment relative to the cytosol. A similar metric was calculated for the single-cell data, where a log_2_ ratio of the mean (normalized and log transformed) gene expression in SR-Parse counts was divided by that in SR-10x counts. Genes with a positive value were more highly expressed in the SR-Parse data relative to SR-10x.

### Parse Biosciences priming strategy separation

To plot the gene coverage for Parse reads originating from either the oligo-dT primers or random hexamer primers, we used the ‘barcode_data.csv’ file produced from split-pipe. The first-round barcodes with an ‘stype’ or ‘R’ are the random hexamer primed reads, whereas the barcodes marked ‘T’ arise from oligo-dT priming. We used the ‘pB’ tag in the Parse BAM files to separate reads from either of the two priming strategies and subsequently plotted their gene coverage profiles.

### Gene detection saturation analysis

To assess gene detection sensitivity as a function of sequencing depth, reads were subsampled within each cell across a range of target depths. For each cell, reads were sampled without replacement at fractions corresponding to depths from 1, 000 to the per-dataset median read count, in steps of 1, 000. The number of unique genes detected was recorded at each depth, and median gene detection across cells was summarized per depth. To extrapolate beyond observed depths, a three-parameter Hill equation was fit to the median gene detection curve for each dataset using nonlinear least squares, with the asymptotic maximum, half-maximum depth, and Hill coefficient as free parameters. Curves were extrapolated to 100, 000 reads per cell.

### MANE Select transcript length assignment

We used biomaRt in R to assign transcript IDs, MANE Select status, and transcript lengths to each gene ID (30, 54). MANE Select defines one transcript at each locus across the genome that is representative of biology at that locus, including protein-coding genes and a targeted subset of lncRNAs and non-coding genes. We retained only the MANE Select transcripts for all genes and these were used to define transcripts length in Figure 2C and B, as well as Supplementary Figure 4B.

### Transcript length distribution analysis

To characterize the distribution of transcript lengths captured by each platform, gene-level counts were weighted by their transcript lengths for each gene and plotted as a kernel density estimate. As a reference for the expected length distribution of polyadenylated transcripts, poly(A)-selected RNA from K562 cells was analyzed on a TapeStation (Agilent). Three technical replicates were prepared from RNA extractions of the same cell population and molarity values were averaged across replicates at each fragment length. The resulting distribution was normalized by area under the curve to place it on a density scale comparable to the sequencing data, and overlaid on the per-platform transcript length distributions. We also used these transcript lengths to compute partial Spearman correlations using the ppcor package in R. This was used to independently assess the contribution of transcript length and nuclear enrichment score to increased SR-Parse expression compared to SR-10x.

### Gene detection sensitivity and biotype analysis

To estimate the practical sensitivity of each platform for detecting individual genes, gene-level counts were summed across all cells and normalized to counts per million (CPM) within each dataset. For genes detected by multiple platforms, CPM values were averaged across detecting methods. Expected counts per cell at a sequencing depth of 50, 000 reads were then derived from the average CPM, and the number of cells required to observe at least one read per gene was estimated as the reciprocal of this value.

Genes were classified by their detection pattern across platforms: detected by all four methods (consensus), detected by three or two methods, or detected exclusively by a single platform. Each gene was assigned a biotype from Gencode v48 annotations and grouped into broad categories: protein-coding, lncRNA, pseudogene, small RNA, mitochondrial, and other. The biotype composition of each detection category was then compared to assess whether platform-specific genes were enriched for particular transcript classes.

### Cross-platform concordance analysis

To assess concordance between the three 10x-based datasets at the per-cell level, UMI and gene counts were compared pairwise across cells with matching barcodes. Per-gene concordance was assessed by comparing log10-transformed total counts across all pairwise dataset combinations, including SR-Parse. For visualization, values were binned into an 80×80 grid and cell density within each bin was plotted, with Pearson’s r and per-axis medians annotated. All concordance analyses were restricted to the set of 20, 164 genes detected across all four datasets.

### Target gene knockdown estimation and differential gene expression

Prior to differential expression analysis, we first created pseudoreplicates by randomly assigning cells into three groups per target gene (55, 56). Each pseudoreplicate was then pseudobulked to create count matrices. We then used DESeq2 to perform the differential expression analysis per target gene (35). Genes with an adjusted p-value < 0.05 were considered statistically significant. To estimate the knockdown of the three target genes, we used the log_2_ fold changes reported by DESeq2. For the power analysis of differential expression between short-read datasets we used powsimR (57).

### Clustering and cell state annotation

SR-10x data was processed using Seurat. Cells were retained if they had a minimum of 3, 000 genes detected, at least 10, 000 UMI counts, and fewer than 10% mitochondrial reads. Data were normalized using log-normalization with a scale factor of 10, 000. The 2, 000 most variable genes were identified using the variance-stabilizing transformation method and used for principal component analysis (PCA). Dimensionality reduction and clustering were performed using the top 30 principal components. A shared nearest-neighbor graph was constructed and Leiden clustering applied at a resolution of 0.4, yielding four clusters. UMAP coordinates were computed using the same 30 components.

Cell states were annotated by scoring each cell against four gene signatures using Seurat’s AddModuleScore: cell cycle activity, erythroid differentiation, stress response, and a low-proliferative state. Signatures were derived from cluster marker genes identified using FindAllMarkers (Wilcoxon rank-sum test, minimum detection fraction 0.25, log_2_FC threshold 0.25). The full gene lists for each signature are provided in Supplementary Table 2.

### Transcriptome-wide impact analysis

To estimate the transcriptome-wide impact of each perturbation independent of significance thresholds, we applied TRADE (Transcriptome-wide Analysis of Differential Expression) to DESeq2 results from the SR-10x dataset using the TRADEtools R package (https://github.com/ajaynadig/TRADEtools) (32). TRADE models the distribution of true effect sizes from observed log_2_ fold changes and their standard errors, providing two summary metrics: transcriptome-wide impact (TI), a measure of the overall perturbation effect across all genes, and the number of effective DEGs (Me), an estimate of the number of genes with non-zero true effects. Analysis was run in univariate mode for each perturbation separately.

### Detection of splicing events with LeafCutter

First, the command ‘regtools junctions extract’ from regtools was used with default parameters to extract junctions from each BAM file for the four datasets (58). The LeafCutter ‘leafcutter_cluster_regtools.py’ script was subsequently run on the junction files for each group to produce the junction cluster count files (33). The script was run with default parameters, which required 50 split reads supporting each cluster. To assess the testable splicing events in each technology, we used these junction cluster counts files.

### Analysis with FLAIR

FLAIR (v3) was used to identify differential gene and isoform expression with DESeq2, and differential isoform usage (DIU) and alternative splicing events (ASEs) with DRIMSeq that were associated with target gene knockdowns (19, 34, 35). Three pseudobulked-pseudoreplicate FASTQ files were provided for the target gene and NTC groups. The FLAIR quantify module estimated transcript abundance by mapping reads to Gencode v48 isoforms, and FLAIR diffsplice and diffexp modules were used to identify DE, DIU and ASEs resulting from the perturbations. We required a difference in isoform fraction (ΔdIF) > 0.1 and an adjusted p-value < 0.05 for significant DIU. Splicing events with ΔPSI > 0.05 and an adjusted p-value < 0.05 were identified as significant ASEs.

To estimate the sequencing depth required for robust DIU detection, target replicate FASTAs were downsampled at 1 million read intervals from 1 million reads up to the maximum available depth, with NTC replicates retained at full depth. Five independent subsampling seeds were used to account for sampling variance. At each depth, isoform abundance was quantified using FLAIR quantify against the Gencode v48 transcriptome, followed by DRIMSeq differential isoform usage testing. Significant DIU isoforms were defined as those with false discovery rate (FDR) < 0.05 and a change in isoform fraction (ΔIF) exceeding the specified threshold. The mean number of significant isoforms across seeds was calculated per depth and target gene. A two-parameter asymptotic exponential model was fit to the mean values for each target and ΔIF threshold, and extrapolated beyond the observed depth range to estimate saturation. Given its pronounced DIU phenotype, *GEMIN5* was used to estimate the depths required to reach 60% and 80% of the asymptotic maximum.

### Protein interaction analysis

Functional consequences of perturbation-induced isoform switches were assessed at the protein level by linking differential isoform usage to predicted changes to protein-protein interactions. Significant DIU events were filtered retaining pairwise isoform shifts between two isoforms that were both protein coding, defined as cases where both the losing and gaining isoforms were annotated as protein-coding in Gencode v48. For each gene, the isoform with the largest decrease in usage (losing isoform) and the isoform with the largest increase in usage (gaining isoform) were selected to represent the dominant switch. Protein sequences for each isoform were retrieved from the Gencode v48 protein translation FASTA.

Known interaction partners for each gene with a protein-coding isoform switch were retrieved from BioGRID (release 4.4.249) and STRING (v12) (37, 38). To focus on high-confidence physical interactions, a three-tier evidence strategy was applied. Tier A interactions required at least two distinct reliable experimental methods in BioGRID (Affinity Capture-MS, Affinity Capture-Western, Reconstituted Complex, Co-fractionation, Proximity Label-MS, or Biochemical Activity). Tier B interactions required one reliable BioGRID method combined with a STRING experimental score ≥ 0.7. Tier C interactions were supported by a STRING experimental score ≥ 0.9 with no BioGRID requirement. Partner protein sequences were retrieved using the MANE Select transcript for each partner gene.

For each protein-coding isoform switch, both the losing and gaining isoform sequences were paired with each interaction partner sequence and submitted to Boltz-2 for protein complex structure prediction (39). Boltz-2 predictions were run with 3 recycling steps, 30 diffusion sampling steps, and 1 diffusion sample per pair. The interface predicted TM-score (ipTM) was used as the primary metric of interaction confidence. For each gene-partner pair, delta ipTM was calculated as the gaining isoform ipTM minus the losing isoform ipTM. To downweight interaction changes at the extremes of the ipTM range, delta ipTM was weighted by a confidence term derived from the mean ipTM of the two isoforms: score = (ipTM_gaining − ipTM_losing) × (4m(1 − m))², where m is the mean ipTM of the two isoforms. This term approaches zero when both isoforms have uniformly low or uniformly high ipTM scores. Pairs with a score ≥ 0.1 were classified as interaction gains and pairs with a score ≤ −0.1 as interaction losses.

### AlphaFold3 structural modeling

To assess the effect of the AK2 isoform switch on AIFM1 binding, we modeled the AK2-214/AIFM1 and AK2-206/AIFM1 complexes using the AlphaFold3 web server (41). Protein sequences were obtained from NCBI CCDS (AK2-214: CCDS374, AK2-206: CCDS81295) and UniProt (AIFM1: O95831), with the N-terminal mitochondrial membrane-binding region of AIFM1 (first 103 amino acids) excluded. Predicted aligned error (PAE) plots and confidence metrics (pTM and ipTM) were used to assess overall fold quality and interface confidence respectively.

## Results

### Experimental design and sequencing platform characterization

We targeted three genes (*DDX6*, *GEMIN5* and *GFI1B*) for knockdown with CRISPRi in the human myelogenous leukemia K562 cell line. These genes have functionally diverse roles in transcriptomic regulation. *DDX6* regulates post-transcriptional gene expression through mRNA degradation (22), *GEMIN5* is an RNA-binding protein and splicing regulator (23), and *GFI1B* is a transcriptional repressor that coordinates hematopoietic gene expression programs (24). We designed guide RNAs (gRNAs) to target the transcription start sites of each gene, as well as five non-targeting control (NTC) gRNAs. Cells were divided into two pools for single-cell library preparation with either 5’ 10x Genomics Chromium (10x) or Parse Biosciences Evercode (Parse). The 10x library was subsequently sequenced using three technologies: Illumina (standard short-read sequencing, SR-10x), PacBio (long-read sequencing, LR-PB), and Oxford Nanopore Technologies (long-read sequencing, LR-ONT). The Parse library was only sequenced with Illumina short-read sequencing (SR-Parse).

After applying quality-control filters to the four datasets and filtering for cells containing only one gRNA (see Methods), we retained 2, 491 cells from the 10x library and 574 cells from the Parse library for analysis (**Supplementary Figure 1A-E**). Read depths varied across datasets, with both long-read methods producing comparable depths per flow cell (**Supplementary Figure 1F**). Given that library preparation influences which genes are captured, we assessed gene detection across platforms at varying read depths. Overall, short-read datasets showed higher sensitivity than their long-read counterparts, with SR-Parse data detecting the most genes (**Figure 1A, Supplementary Figure 2A**). Using all available reads, a set of 20, 164 consensus genes were detected across all four datasets. These showed substantially higher expression than genes detected by only a subset of platforms (**Supplementary Figure 2B-C**). Parse identified the largest number of platform-unique genes (n=10, 336), though all datasets detected unique genes (**Supplementary Figure 2B**). Sensitivity is often emphasized as a key differentiator for single-cell platforms (25, 26). To assess the practical value of these method-unique genes, we estimated the number of cells required to detect at least one read per gene at a depth of 50, 000 reads per cell. While consensus genes required a median of 3 cells, dataset-unique genes required a median of 334–709 cells depending on platform (**Supplementary Figure 2D**). Platform-unique genes were also enriched for lncRNAs and pseudogenes, biotypes conventionally associated with low expression levels (**Supplementary Figure 2E**). Taken together, these results caution against equating higher gene counts with improved sensitivity to functionally meaningful genes.

**Figure 1.**
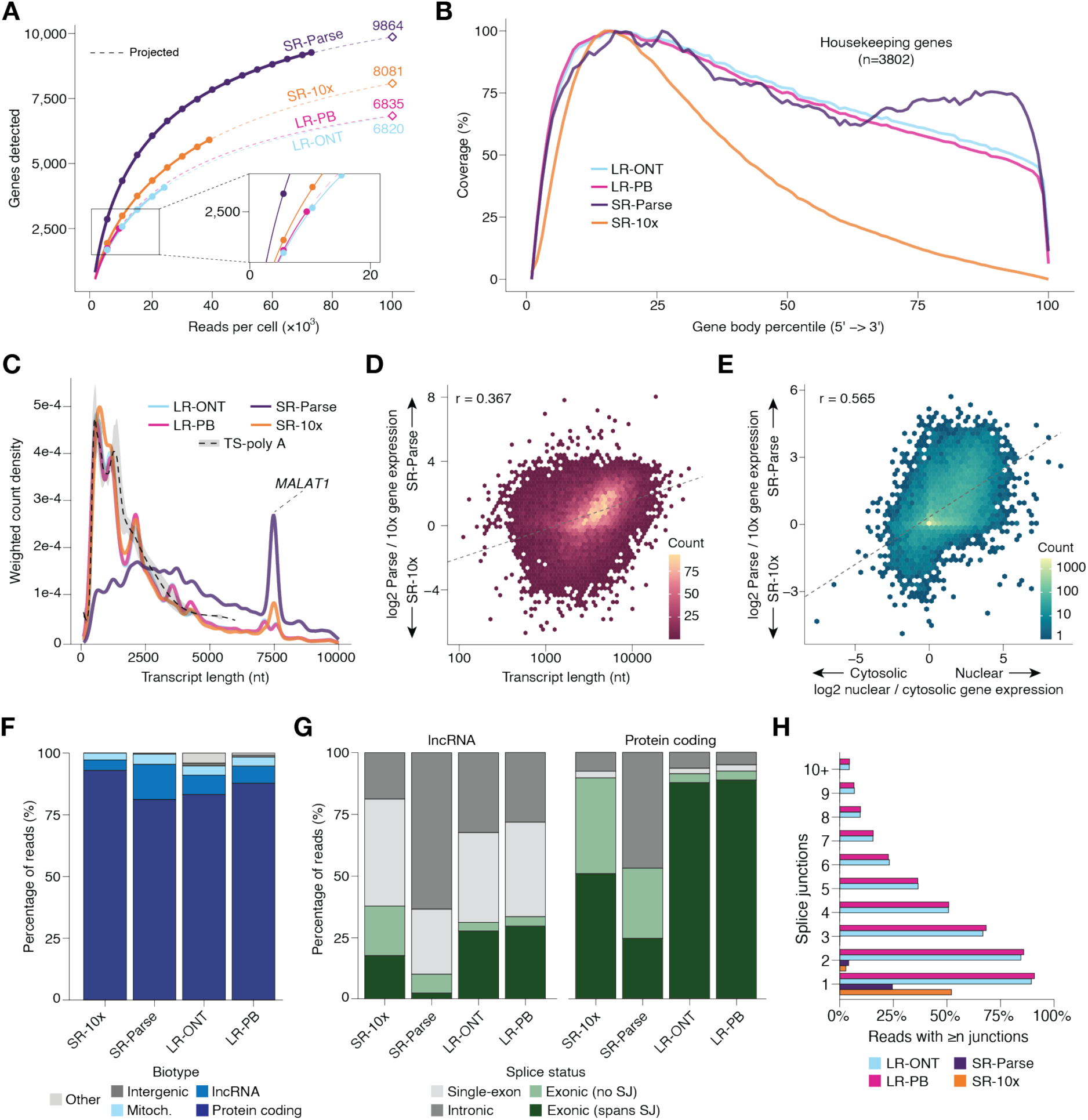
Characterization of detected genes across sequencing platforms. **A.** Gene detection per method as a function of sequencing depth per cell. Solid lines represent observed data; dashed lines are extrapolated using the Hill equation. **B.** Normalized gene body coverage per method across 3, 802 housekeeping genes was plotted using RSeQC (27). **C.** Transcript length distributions weighted by gene counts per dataset. The ‘MANE Select’ transcripts were chosen for each gene. Dashed line: poly(A)-selected RNA fragments from K562 cells (TapeStation). The large peak represents transcripts mapped to the lncRNA gene *MALAT1*. **D.** A log_2_ ratio was calculated of the mean (normalized and log transformed) gene expression in SR-Parse counts divided by SR-10x counts. Genes with a positive value are more highly expressed in the SR-Parse data relative to SR-10x. This ratio was plotted against the MANE Select transcript length for each gene. Points are colored by count density. Spearman ρ=0.394, p<0.001. **E.** The y-axis values were calculated as in panel D. Public ENCODE gene expression data was used to calculate a log_2_ ratio comparing nuclear and cytosolic expression. Points are colored by log10 transformed count density. Spearman ρ=0.565, p<0.001. **F.** Percentage of reads mapped to different gene transcript biotypes. **G.** The splice status of reads mapped to lncRNA (left) and protein-coding (right) transcripts. lncRNA: long non-coding RNA; Mitoch.: mitochondrial; SJ: splice junction. **H.** Percentage of reads spanning one or more splice junctions per dataset.

As uniform coverage across transcripts is a prerequisite for comprehensive splice junction detection, we assessed gene body coverage profiles using a set of 3, 802 previously established housekeeping genes (27) (**Figure 1B**). SR-10x displayed a strong 5’ bias, consistent with its 5’-capture design. SR-Parse showed more evenly distributed coverage across gene bodies, reflecting its use of random hexamer priming alongside oligo-dT. Both long-read datasets showed more uniform coverage than SR-10x, but 5’ bias was still evident. This likely reflects internal oligo-dT priming, where primers bind adenine-rich sequences within transcripts rather than exclusively at poly(A) tails, resulting in truncated capture and under-representation of 3’ regions (28, 29). Despite these coverage differences, gene quantification was highly consistent across sequencing approaches when library preparation was held constant. Per-cell counts and gene detection were highly correlated across the three 10x-based datasets, which represent the same cells sequenced with different technologies (r > 0.90; **Supplementary Figure 3A-B**). Gene-level expression profiles showed the same pattern, 10x datasets were highly concordant (r > 0.88), while SR-Parse showed weaker correlation with all other platforms (r = 0.52–0.62; **Supplementary Figure 3C**). This suggests that even across fundamentally different sequencing technologies, library preparation was the stronger determinant of expression profiles.

To further assess patterns introduced by library preparation approaches, we examined the transcript length distribution captured by each method. For each dataset, detected genes were assigned their MANE Select transcript length which we plotted against the weighted gene counts (30). To establish a reference for the expected distribution of polyadenylated transcripts, we included an RNA fragment distribution from a TapeStation analysis of poly(A)-selected transcripts in K562 cells (see Methods) (**Figure 1C**). All three 10x-based datasets showed largely similar profiles, closely mirroring the mRNA fragment size distribution. In contrast, Parse showed a profile skewed toward longer transcripts (**Figure 1C-D**).

Two important features that distinguish Parse from 10x library preparation are random hexamer priming and cell fixation with permeabilization. To investigate which drives the preferential capture of longer transcripts, we first compared random hexamer and oligo-dT-primed reads within the SR-Parse library. Random hexamer-primed reads showed a modest enrichment for longer transcripts compared to oligo-dT-primed reads (**Supplementary Figure 4A**), suggesting hexamer priming contributes to, but does not fully explain, the observed length bias.

We therefore investigated whether fixation and permeabilization leads to preferential capture of nuclear RNA. Comparing per-gene SR-Parse expression relative to SR-10x against nuclear enrichment scores derived from publicly available K562 fractionation data (see Methods), we found a strong positive correlation between Parse enrichment and nuclear localization (Spearman ρ=0.56, p<2.2×10^-16^, **Figure 1E**). As transcript length and nuclear enrichment score were also found to be moderately correlated (Spearman ρ=0.41, p<2.2×10^-16^), we computed partial Spearman correlations to assess the independent contribution of each. Both nuclear enrichment (partial ρ=0.48, p<2.2×10^-16^) and transcript length (partial ρ=0.39, p<2.2×10^-16^) independently predicted increased Parse expression after controlling for the other. As nuclear transcripts are substantially longer than cytosolic transcripts (Mann-Whitney U p<2.2×10^-16^, **Supplementary Figure 4B**), these results indicate that nuclear RNA capture is the primary driver of the length bias, with hexamer priming contributing a secondary effect.

Consistent with nuclear RNA enrichment, Parse showed a higher fraction of lncRNAs, which are predominantly nuclear-localized (31), and a correspondingly lower fraction of protein-coding transcripts (**Figure 1F**). A high degree of intronic reads reinforced this pattern. Nearly half of Parse protein-coding reads and over 60% of lncRNA reads were intronic, a hallmark of nuclear RNA capture (5, 26), far exceeding the fraction of intronic reads captured in other datasets (**Figure 1G**). To assess implications for splicing analysis, we quantified how many reads contained a splice junction between the different datasets. The high intronic content of Parse libraries limited splice junction capture, with fewer than 25% of reads spanning a single splice junction, compared to approximately 50% for SR-10x and over 80% for long-read datasets **(Figure 1H**).

Taken together, these results demonstrate that library preparation is a stronger driver of variation between datasets than sequencing approach. Parse’s use of random hexamer primers provides more even gene body coverage, but permeabilization leads to nuclear enrichment, resulting in a high proportion of intronic reads, limiting splice junction capture. Meanwhile, long-read datasets showed the highest proportion of junction-spanning reads, despite lower overall sensitivity. These findings highlight that platform suitability depends on the biological question: gene expression versus isoform resolution requires different approaches.

### Assessing gene-level responses to CRISPRi knockdown

We next examined the transcriptional consequences of each perturbation. To assess knock-down efficiency, we performed a differential expression (DE) analysis between perturbed and non-targeting control (NTC) cells using pseudobulked counts based on replicates from randomly assigned cells (see Methods, **Supplementary Table 1**). Guide RNAs that did not result in knockdown of their target gene were excluded from downstream analysis (log_2_ fold change > –0.5). Remaining gRNAs showed consistent knockdown of their target genes across all four platforms (**Supplementary Figure 5A-C**). Dimensionality reduction of SR-10x data revealed four major clusters (**Figure 2A**), corresponding to cell cycle activity, erythroid differentiation, stress response, and a low-proliferative state (**Figure 2B**, see Methods, **Supplementary Table 2**). *DDX6* and *GFI1B* knockdown cells were enriched in the low-proliferative cluster relative to non-targeting controls, with *DDX6* also enriched in the stress cluster (**Figure 2C, Supplementary Figure 6A**), consistent with knockdown of essential genes reducing proliferative capacity (4).

**Figure 2.**
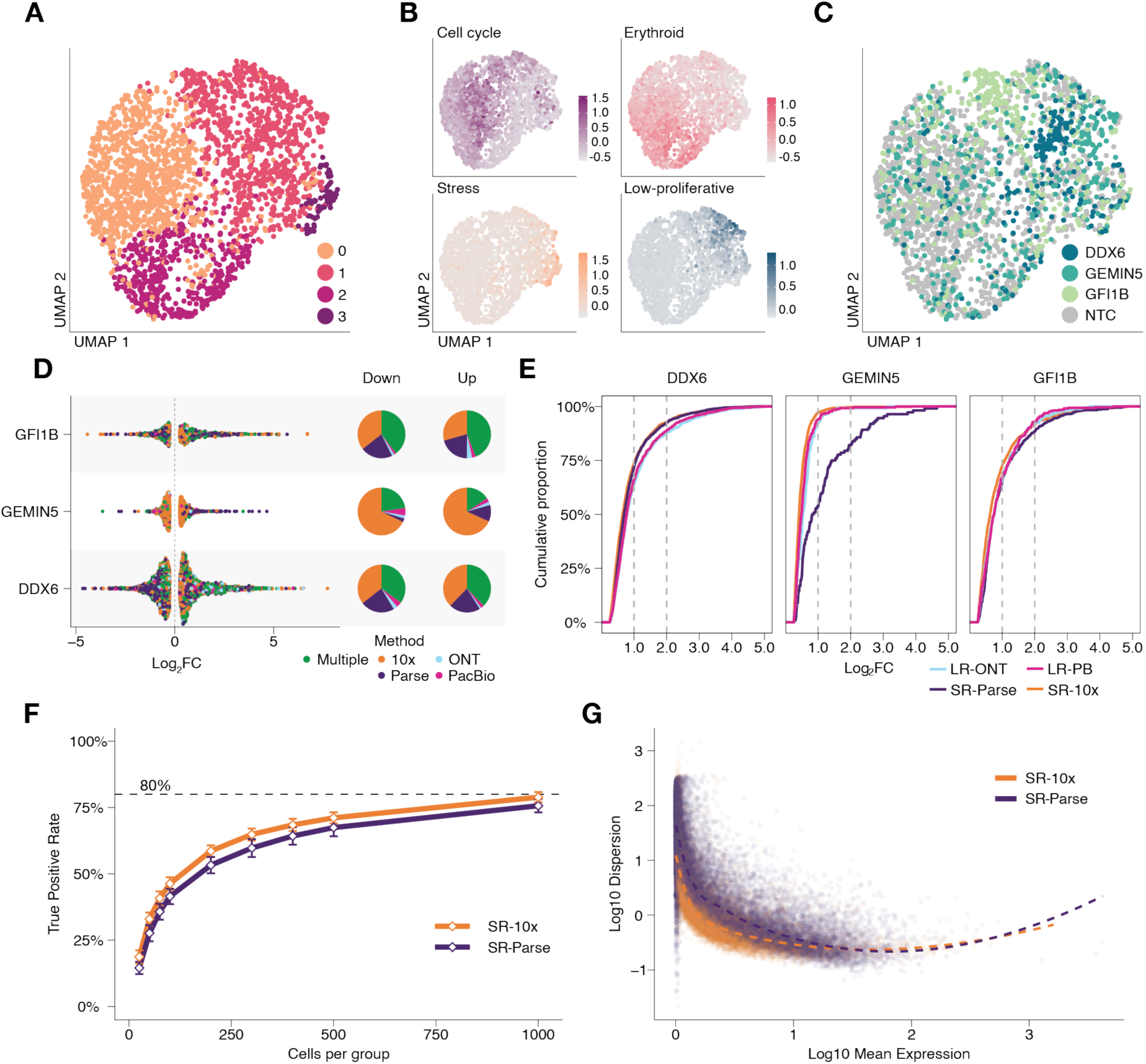
Gene-level transcriptional responses to CRISPRi knockdown across platforms. **A.** UMAP of SR-10x cells (n=2, 491) colored by Seurat cluster identity (0–3). **B.** UMAPs as in (A), colored by module scores for four annotated cell states: cell cycle activity, erythroid differentiation, stress response, and low-proliferative state (see Methods, Supplementary Table 2). **C.** UMAP as in (A), colored by perturbation identity. **D.** Left: log_2_ fold change (FC) of DEGs (DESeq2, |log_2_FC| > 0.25, false discovery rate (FDR) < 0.05) per perturbation, colored by the platform(s) in which each gene was detected as a DEG. Right: Pie charts showing the proportion of down- and upregulated DEGs detected per platform. **E.** Cumulative proportion of DEGs per perturbation and platform as a function of absolute log_2_FC. Dashed vertical lines indicate |log_2_FC| = 1 and 2. **F.** True positive rate across simulated cell numbers for SR-10x and SR-Parse (powsimR); effect sizes sampled from a gamma distribution. Dashed line: 80% recall rate. **G.** Mean–dispersion relationship for SR-10x and SR-Parse; dashed lines indicate fitted trends.

Analyzing the transcriptome-wide effects of each perturbation, *DDX6* knockdown produced the strongest differential expression signal across all platforms, while *GEMIN5* produced the weakest (**Figure 2D-E**, **Supplementary Figure 6B**). A substantial fraction of differentially expressed genes were detected by multiple platforms, though each method identified some unique hits (**Figure 2D, Supplementary Table 3**). Effect size distributions were comparable across platforms within each perturbation, with *DDX6* and *GFI1B* showing a larger proportion of genes with strong effects (|log_2_FC| > 1) than *GEMIN5* (**Figure 2E**). Notably, SR-Parse detected very few DEGs for *GEMIN5* knockdown relative to other platforms.

To assess whether the weak differential expression signal of GEMIN5 reflects a genuinely limited transcriptional response or simply small per-gene effects spread across many targets, we applied TRADE (Transcriptome-wide Analysis of Differential Expression) (32). TRADE estimates transcriptome-wide impact independent of significance thresholds by modeling the distribution of true effect sizes. This analysis confirmed that *DDX6* knockdown produced the largest transcriptome-wide impact, affecting approximately 3, 000 genes, and revealed that *GEMIN5* affected a large number of genes, but with small per-gene effects (**Supplementary Figure 6C-D**). This pattern is consistent with *GEMIN5*’s role as a splicing regulator, where phenotypic consequences may manifest at the isoform level rather than through changes in total gene expression.

Overall, short-read datasets identified more differentially expressed genes than long-read datasets (**Supplementary Figure 6B**), likely reflecting differences in sequencing depth and cell numbers rather than platform sensitivity. To directly compare DE detection power between the two short-read platforms independent of these confounders, we used powsimR [21], which estimates the mean-dispersion relationship from each dataset and simulates counts to assess statistical power at matched sample sizes. At matched sample sizes, SR-10x showed higher true positive rates for DE detection than SR-Parse (**Figure 2F**), indicating that 10x libraries provide greater statistical power per cell. This difference is explained by higher gene-level dispersion in SR-Parse across all expression levels (**Figure 2G**), indicating greater technical noise.

Despite these differences in detection power, the platforms showed strong agreement in estimated effect sizes for genes detected as differentially expressed. Log_2_ fold change estimates were well-correlated across all four platforms (**Supplementary Figure 7A**), with the highest concordance between the two long-read methods (Spearman ρ=0.91, p<0.001; **Supplementary Figure 7B-C**). This indicates that for gene-level analysis, all platforms reliably capture major perturbation effects, though they vary in the sequencing depth required, and therefore cost, to achieve equal sensitivity varies.

### Comparison of alternative splicing event detection across platforms

We next examined the splicing consequences of each perturbation. We used LeafCutter to compare splice junction quantification across all datasets, as it does not require isoform inference and can be applied to both short- and long-read data (33). Across the 10x-based datasets, all three methods captured thousands of splice junctions with sufficient coverage for testing (**Figure 3A**, **Supplementary Table 4**), while SR-Parse yielded far fewer, consistent with its high intronic read content. Despite lower overall read depth, both long-read datasets captured more testable splice junctions than either short-read method, with LR-PB yielding the most overall. The distribution of testable alternative splicing events (ASEs) across gene bodies varied markedly across platforms (**Figure 3B**). SR-10x showed the strongest 5’ enrichment, with an approximately 8-fold difference in testable splice junction density between the 5’ and 3’ ends of transcripts. The LR datasets showed a more moderate but still evident 5’ bias, mirroring the read coverage profiles observed above (**Figure 1B)**. Surprisingly, despite the incorporation of random hexamers during reverse transcription, SR-Parse also exhibited a 5’ bias. Importantly, this platform-dependent positional skew has practical consequences for splicing analysis. Stratification by transcript length revealed that the 5’ bias in SR-10x becomes progressively more severe for longer transcripts, while it remains less pronounced in the LR datasets and SR-Parse (**Supplementary Figure 8A**), meaning that splicing results from SR-10x data will be increasingly incomplete for longer genes.

**Figure 3.**
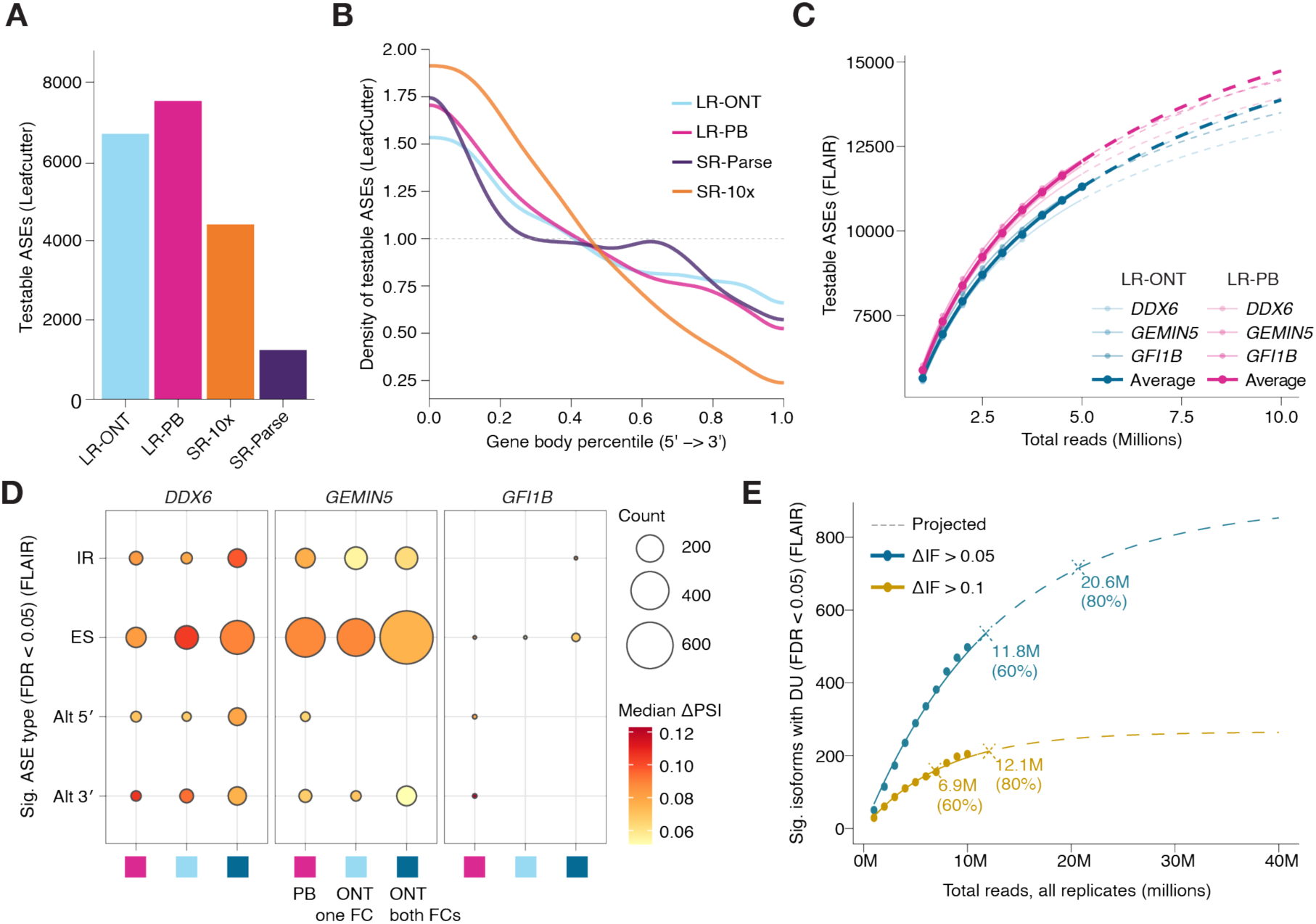
Splicing detection sensitivity varies across platforms and sequencing depths. **A.** Total testable alternative splicing events (ASEs) detected by LeafCutter per dataset. **B.** Density of testable ASE positions from LeafCutter plotted along normalized gene bodies for each dataset. **C.** Testable ASEs from FLAIR across read depths, comparing one flow cell per platform. Individual perturbations are shown as transparent lines; opaque lines indicate the mean across all perturbations. **D.** Significant ASEs from FLAIR (ΔPSI (percent spliced in) > 0.05, FDR < 0.05) per perturbation and dataset. Circle size: number of events; color: median ΔPSI. IR: intron retention; ES: exon skipping; Alt 5’/3’: alternative 5’/3’ splice site usage. Colored boxes on x-axis represent long-read sequencing methods, PacBio (PB), Oxford Nanopore Technologies (ONT) with one flow cell (FC) or both flow cells. **E.** Isoforms with significant differential usage (ΔIF (isoform fraction) > 0.05 or ΔIF > 0.1, FDR < 0.05) in *GEMIN5* knockdown cells from FLAIR, calculated across read depths using both LR-ONT flow cells. Dashed lines indicate projected saturation curves.

Comparing a single flow cell per platform, LR-PB identified more testable ASEs than LR-ONT across all read depths (**Figure 3A**). However, as LeafCutter was designed for short-read data and its junction detection is sensitive to sequencing error rates (13), the higher error rate of ONT reads may partially explain this difference. To assess whether this advantage holds when using a tool designed for long-read data, we applied FLAIR, which explicitly accounts for higher error rates in long-read sequencing, to identify differential isoform expression (DIE), differential isoform usage (DIU), and ASEs in both LR datasets (**Supplementary Table 5**) (19, 34, 35). LR-PB identified more testable ASEs than LR-ONT across all read depths (**Figure 3C**), suggesting that PacBio’s accuracy advantage is not an artifact of the LeafCutter analysis but reflects a genuine sensitivity difference for splicing detection.

To assess statistical power across perturbations and sequencing depths, we incorporated a second ONT flow cell, adding 119 million reads for a total of 232 million ONT long-reads. Across perturbations, *GEMIN5* knockdown produced substantially more significant ASEs than *DDX6*, with exon skipping as the dominant event type, while *GFI1B* produced too few events for meaningful comparison (**Figure 3D**, **Supplementary Figure 8B**). This splicing phenotype stands in stark contrast to the gene-level analysis, where *GEMIN5* showed the weakest differential expression signal of the three perturbations (**Figure 2D-E**). Given its pronounced splicing phenotype, we used *GEMIN5* to estimate the sequencing depth required for robust isoform-level detection (**Figure 3E**). Using differential isoform usage as the readout, at a change in isoform fraction (ΔIF) threshold of 0.05 approximately 20.6 million reads were required to reach 80% saturation, while a less stringent threshold of ΔIF > 0.1 required approximately 12.1 million reads.

### Long-read perturbation screens reveal isoform-level effects with distinct protein-interaction consequences

Knockdown of *GEMIN5* produced the most extensive splicing changes of the three perturbations, yet the weakest gene-level signal. We therefore chose to further examine how perturbation effects manifest at the isoform level. Using the both flow cells of the ONT dataset, we identified 969 cases of differential isoform usage (DIU) within 616 genes (**Figure 4A**). We assessed the relationship between differential gene expression and DIU across all three perturbations (**Figure 4B**). Strikingly, the three perturbations showed markedly distinct patterns. For *GFI1B*, isoform expression changes were tightly coupled to gene expression changes (r = 0.88), suggesting that the isoform-level changes observed are largely driven by overall transcript abundance rather than independent splicing regulation (**Supplementary Figure 8B**). In contrast, *GEMIN5* and *DDX6* knockdown drove extensive DIU largely independent of gene expression changes (r = 0.57 and 0.75 respectively), with large shifts in isoform proportion occurring in genes showing little to no differential expression. These results indicate that for many genes, conventional methods miss a substantial proportion of perturbation effects, which manifest through altered isoform usage rather than changes in overall gene expression.

**Figure 4.**
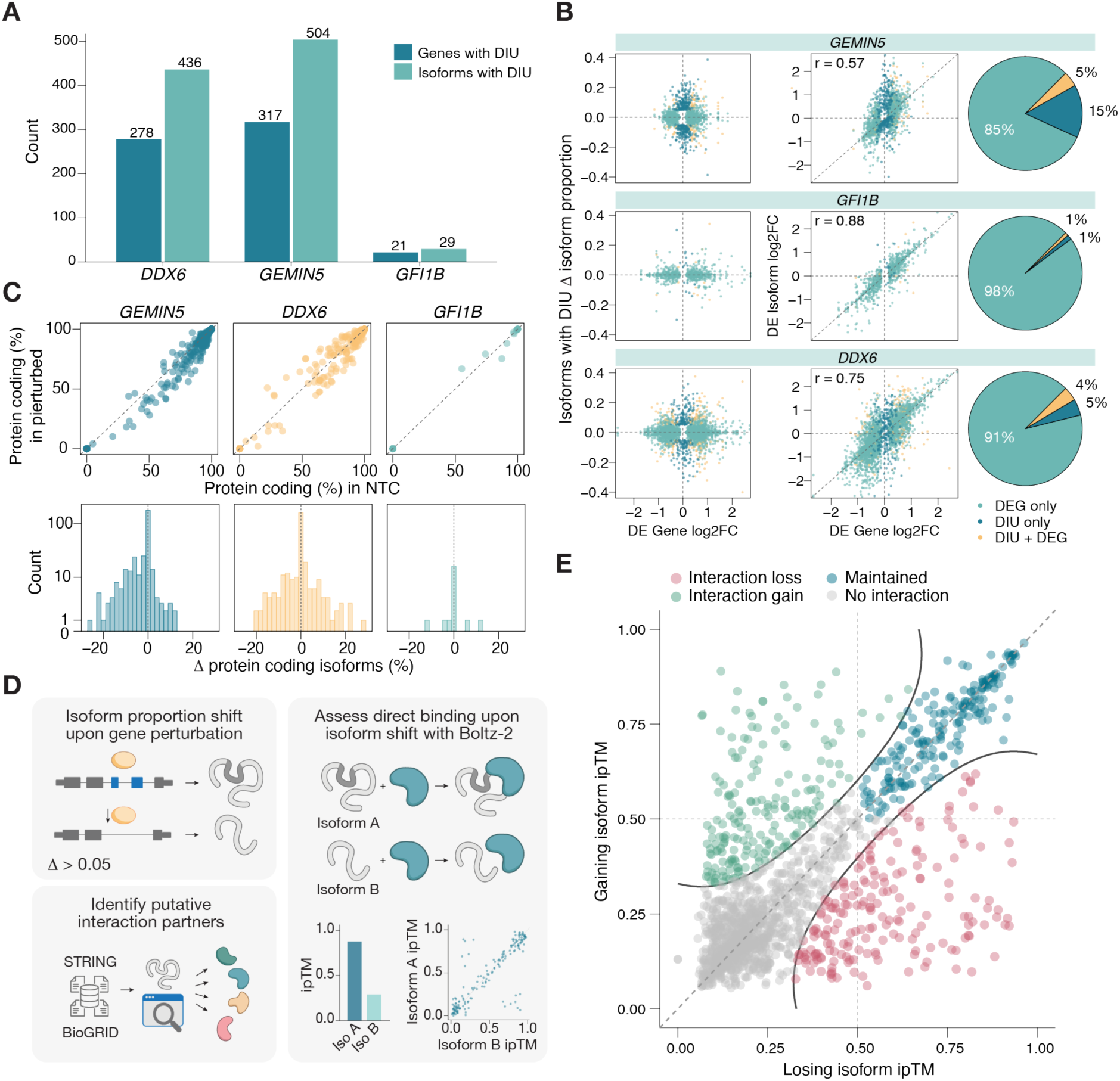
Isoform-level changes result in distinct protein-interaction consequences. **A.** The number of genes containing differential isoform usage (DIU) (dark blue) and isoform with differential usage (light blue) per perturbation identified with FLAIR (change in isoform proportion > 0.05 and FDR < 0.05). **B.** Gene-level versus isoform-level changes per perturbation. Left: change in isoform proportion vs gene log_2_FC; middle: isoform log_2_FC vs gene log_2_FC (Pearson’s r). Points colored by significance: DEG only, DIU only, or both (FDR < 0.05, DRIMSeq). Right: proportion of each category. C. For genes containing DIU, the percentage of protein coding isoforms in perturbed groups is compared to the percentage in the NTC group (top), the change in proportion of protein coding isoforms (bottom) per perturbation is shown. D. Schematic illustrating how changes between protein-coding isoforms alter protein interaction partners. ipTM: interface predicted template modelling score. E. The ipTM scores for DIU gaining (increasing in proportion) isoforms compared with losing (decreasing in proportion) isoforms for all perturbations. Dots are coloured by whether there is an interaction partner gain (green), the interaction is maintained between isoforms (blue), no interaction identified with either isoform (grey), or an interaction partner is lost (red).

To further characterise how perturbation effects impact transcript function, we examined biotype shifts resulting from DIU. A functional asymmetry across perturbations was evident, with *GEMIN5* knockdown producing a systematic shift away from protein-coding isoforms and a corresponding increase in transcripts subject to nonsense-mediated decay and intron retention (**Figure 4C**, **Supplementary Figure 9A**). This pattern is consistent with the role of *GEMIN5* in SMN complex assembly and spliceosome biogenesis (36), and stands in contrast to *DDX6* knockdown, which showed a more balanced redistribution of isoform proportions with no systematic directional shift.

Finally, we sought to assess the functional consequences of perturbation-induced isoform shifts at the protein level. Recent advances in protein structure prediction have enabled modeling of protein-protein interactions at scale, and we developed a pipeline leveraging these approaches to estimate how isoform switches affect the ability of proteins to interact with their known binding partners (**Figure 4D**, Methods). For each gene with a switch between protein-coding isoforms, we identified putative interaction partners from BioGRID and STRING, and used Boltz-2, a scalable open-weight protein foundation model, to predict complex structures for both the losing and gaining isoforms paired with each partner (**Supplementary Figure 10A-D**) (37–39). Using this approach we predicted interaction confidence for 3, 414 isoform-partner pairs across 116 genes with protein-coding isoform switches, yielding 1, 650 complete pairwise comparisons. Comparing the interface predicted TM-score (ipTM) between losing and gaining isoforms, 391 pairs showed substantial changes in predicted interaction confidence, comprising 197 interaction gains and 194 interaction losses (**Figure 4E**).

### Isoform-resolution analysis reveals functional consequences of GEMIN5 perturbation

To demonstrate how isoform-level analysis can reveal the precise functional consequences of a perturbation, we examined *AK2*, which encodes adenylate kinase 2, a protein that maintains adenine nucleotide balance in the mitochondrial intermembrane space, a function essential for ADP/ATP carrier activity and cellular respiration. AK2 exerts this function through interaction with AIFM1 on the mitochondrial membrane, an interaction recently shown to be mediated by a C-terminal domain that anchors AK2 to its partner (**Supplementary Figure 11A**) (40). Following *GEMIN5* knockdown, *AK2* showed significant upregulation at the gene level (**Figure 5A**). This increase was accompanied by a substantial shift in isoform proportions away from the canonical AK2-214 isoform, and towards AK2-206, which lacks exon 6. No changes in *AK2* expression or isoform usage were observed following *DDX6* or *GFI1B* knockdown, indicating that these effects are directly mediated by *GEMIN5* perturbation (**Supplementary Figure 11B-C**). To assess how this isoform shift affects AK2–AIFM1 interactions, we used AlphaFold3 to predict the binding interface with and without the exon 6-encoded domain. While modeling AK2-214/AIFM1 recapitulated the C-terminal binding interface previously described (**Figure 5B**), AK2-206/AIFM1 modeling instead predicted binding on the opposite face of AIFM1, with interaction confidence dropping sharply (ipTM: 0.59 → 0.21; **Figure 5C–D**, **Supplementary Figure 11D**) (41).

**Figure 5.**
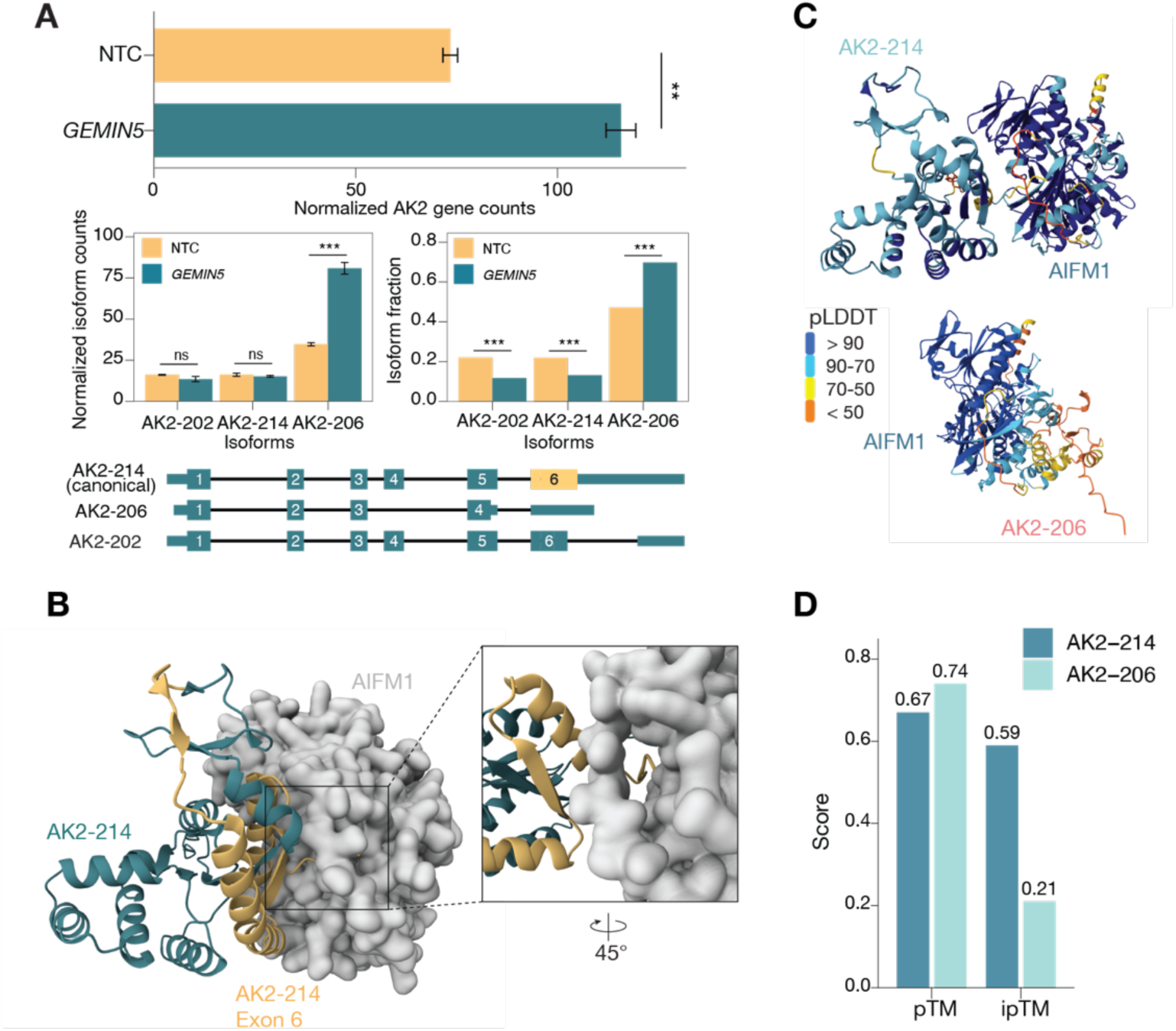
Isoform-level analysis reveals perturbation effects independent of gene expression changes upon *GEMIN5* knockdown. **A.** Differential isoform usage analysis of the *AK2* gene upon *GEMIN5* knockdown using FLAIR. Gene expression of *AK2* is shown between non-targeting control cells (NTC) and *GEMIN5* knockdown (top). The isoform expression (middle left) and isoform proportions (middle right) of three *AK2* isoforms are shown. The ** and *** indicate adjusted p-values <0.01 and <0.001 respectively from DESeq2 for differential expression panels and from DRIMSeq for the isoform fraction panel. ns=non-significant. The structures of these three isoforms are displayed (bottom), with boxes representing exons (numbered) and lines representing introns. Thick boxes show the open reading frame of encoded proteins. **B.** AlphaFold3-predicted structure of AK2-214 (canonical) in complex with AIFM1. Exon 6-encoded region highlighted (yellow). Inset: 45° rotation of binding interface. **C.** AlphaFold3-predicted complexes colored by pLDDT confidence score pLDDT: (predicted local distance difference test). Top: AK2-214/AIFM1. Bottom: AK2-206/AIFM1. **D.** AlphaFold3 confidence metrics for predicted AK2/AIFM1 complexes. pTM: predicted template modelling score, ipTM: interface predicted score.

The *AK2* example illustrates how integrating isoform-level analysis with protein interaction modeling can generate precise functional hypotheses from perturbation data. Gene-level analysis alone would attribute *AK2* upregulation to increased expression of AK2-214, the canonical isoform, suggesting enhanced mitochondrial metabolism. Instead, isoform-level analysis reveals that upregulation is driven entirely by AK2-206, and interaction modeling predicts this isoform is incapable of AIFM1 binding. Since AK2 localization to the mitochondrial intermembrane space depends on its interaction with AIFM1, this suggests that the net effect of *GEMIN5* perturbation is not enhanced mitochondrial AK2 activity, but rather its displacement from the mitochondrial membrane. More broadly, this pipeline — from perturbation to isoform switch to predicted protein interaction consequences — provides a framework for generating mechanistic hypotheses that cannot be produced with gene-level quantification alone.

## Discussion

The human genome contains approximately 20, 000 protein-coding genes, but estimates suggest as many as 170, 000 distinct protein-coding transcripts may be expressed (42). Yet, the most widely used single-cell sequencing methods can only resolve gene expression (8, 43). Our platform comparison revealed that at the gene level, library preparation is a stronger driver of transcriptomic variation than sequencing technology, but for isoform-level analysis, both dimensions matter. As most single-cell sequencing approaches have been designed to maximize scale, design choices have also shaped which transcripts are captured. For instance, the cell permeabilization and random hexamer priming employed by Parse Evercode leads to nuclear RNA enrichment and a high proportion of intronic reads, limiting splice junction detection regardless of sequencing platform. These findings underscore that platform choice for isoform-level analysis cannot be based on sequencing technology alone. The biological questions that can be addressed are ultimately determined by the sensitivity and biases inherent to the chosen library preparation and sequencing platform.

This has particular consequences for CRISPR screens, where the goal is not merely to characterize transcriptomes but to detect the functional consequences of perturbations. As CRISPR screens have grown in both number and scale, the vast majority have been performed at gene-level resolution (1, 2, 4, 44), meaning genes with primarily post-transcriptional functions risk having their effects underestimated or missed entirely. This gap in resolution represents a systematic blind spot in functional genomics. Our finding that *GEMIN5* knockdown produces minimal gene-level signal but pervasive differential splicing illustrates this problem; in a conventional screen, *GEMIN5* would appear as a weak hit, when in fact it profoundly reshapes the transcriptome. More concerning, gene-level analysis overlooks isoform-level complexities that can lead to incomplete or misleading biological interpretation, as demonstrated with *AK2* in our dataset. Whether similar interpretive errors exist in cell atlases and disease signatures, built almost exclusively from gene-level data, remains to be explored (45, 46).

Our comparison of four platforms for pooled single-cell CRISPR screens reveals that long-read sequencing uniquely enables isoform-level resolution in perturbation screens. We assess how library preparation methods, sequencing platforms, and analytical approaches affect output, offering practical guidance for researchers designing studies where splicing is central to the biological question. Critically, the use of CRISPR perturbations adds a dimension absent from most benchmarking studies. Rather than evaluating platforms on technical metrics alone, we can directly assess each platform’s ability to detect biologically meaningful changes. Short-read methods cannot resolve full-length isoforms regardless of sequencing depth (10, 11), and as perturbation screens continue to grow in scale, the biology missed at gene-level resolution will represent an increasingly significant blind spot. While current long-read throughput means that screens with hundreds of perturbations remain costly, the landscape is shifting rapidly. Reusable flow cell chemistries, decreasing per-base sequencing costs, and emerging library preparation strategies compatible with scalable single-cell long-read capture collectively suggest that the throughput barrier is close to being overcome (47, 48). Beyond sequencing, the emergence of scalable protein structure prediction models now enables a layer of functional interpretation that was previously out of reach. The framework we present here, linking perturbation induced isoform switches to predicted protein-level consequences, provides a blueprint for how perturbation screens can move beyond transcript quantification towards a mechanistic understanding of gene function.

## Declarations

## Data Availability

The raw FASTQ files and processed single-cell data generated in this study is available through GEO under accession GSE330222.

## Acknowledgements

The authors acknowledge support from the National Genomics Infrastructure in Stockholm funded by Science for Life Laboratory, the Knut and Alice Wallenberg Foundation and the Swedish Research Council, and SNIC/NAISS/Uppsala Multidisciplinary Center for Advanced Computational Science for assistance with massively parallel sequencing and access to the UPPMAX computational infrastructure.

## Author Contributions

T.L., J.G. and N.A. conceived of the study. J.P. performed the experiments, with assistance from S.Ö. and S.L.. J.G. and N.A. performed all computational analyses and created the figures. J.G., N.A. and T.L. wrote the manuscript with contributions and review from all the authors.

