## Supplementary Figures for "Isoform-level resolution in single-cell CRISPR screens reveals hidden functional consequences of gene perturbation"

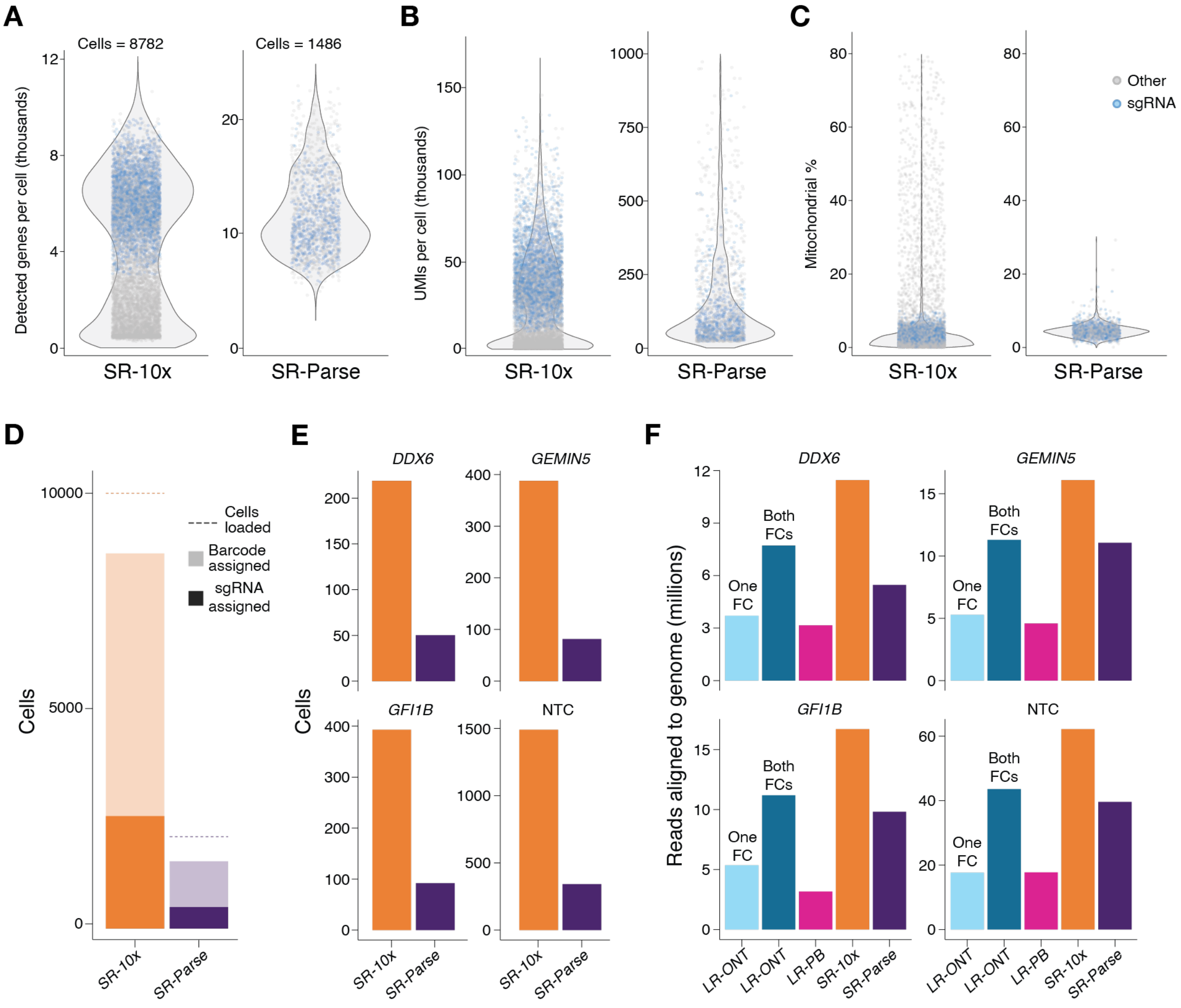


**Supplementary Figure 1. Single-cell quality control of different methods.**

**A.** The number of detected genes (x1000), **B.** Unique molecular identifiers (UMIs) (x1000), and **C.** percentage of mitochondrial genes per cell are plotted for short-read 10x and Parse datasets. Cells that contained a single guide RNA (sgRNA) and were kept for downstream analysis are colored blue. **D.** Bar charts showing the total number of cells kept for analysis in both short-read 10x and Parse datasets. Dashed lines represent the total number of cells loaded per experiment. Light colored bars represent the cells which had a valid cell barcode and dark colored boxes represent the cells which passed QC and had a sgRNA which were kept for analysis. **E.** Bar charts showing the number of cells containing sgRNAs targeting three genes, or non-targeting control sgRNAs for 10x short-read (orange) and Parse short-read (purple). **F.** The number of reads aligned to the genome (in millions) for all datasets across cells containing either non-targeting control or gene-targeting sgRNAs. For the long-read Oxford Nanopore Technologies data (LR-ONT), bars are shown for a single flow cell (one FC) or both flow cells combined (both FCs).

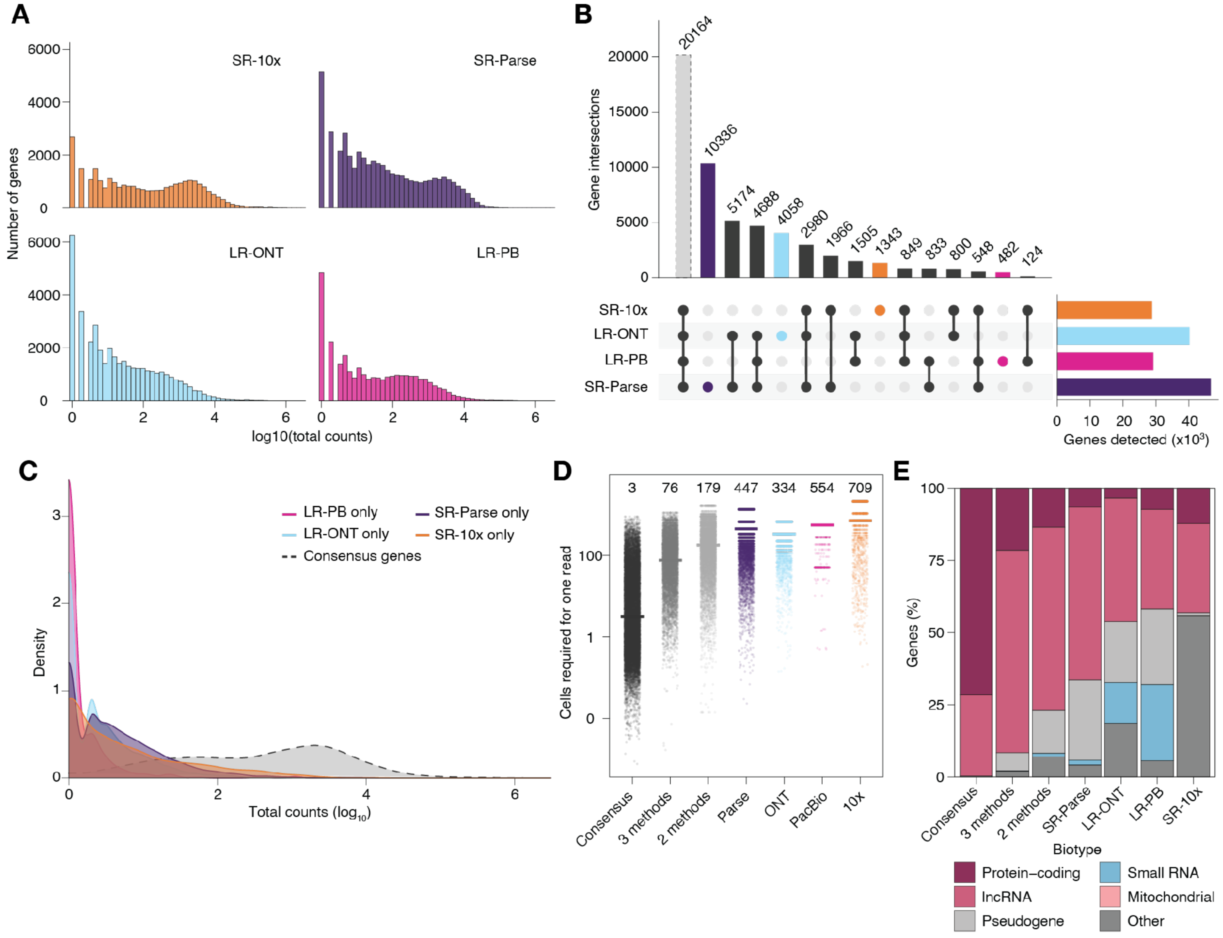


**Supplementary Figure 2. Gene detection across datasets.**

**A.** Distribution of log10-transformed total counts per gene (summed across all cells) for each dataset. **B.** UpSet plot showing the overlap in detected genes across datasets. Bars indicate the number of genes in each intersection; horizontal bars on the right show total genes detected per dataset. **C.** Density of log10-transformed total counts for platform-specific genes and consensus genes (dashed line); consensus genes detected across all four datasets (n=20,164). **D.** Estimated number of cells required to observe at least one read per gene at 50,000 reads per cell, shown on a log10 scale. Numbers above each column indicate the median. Genes are grouped by detection category: consensus, detected by three or two methods, or detected exclusively by a single platform. **E.** Biotype composition of genes in each detection category.

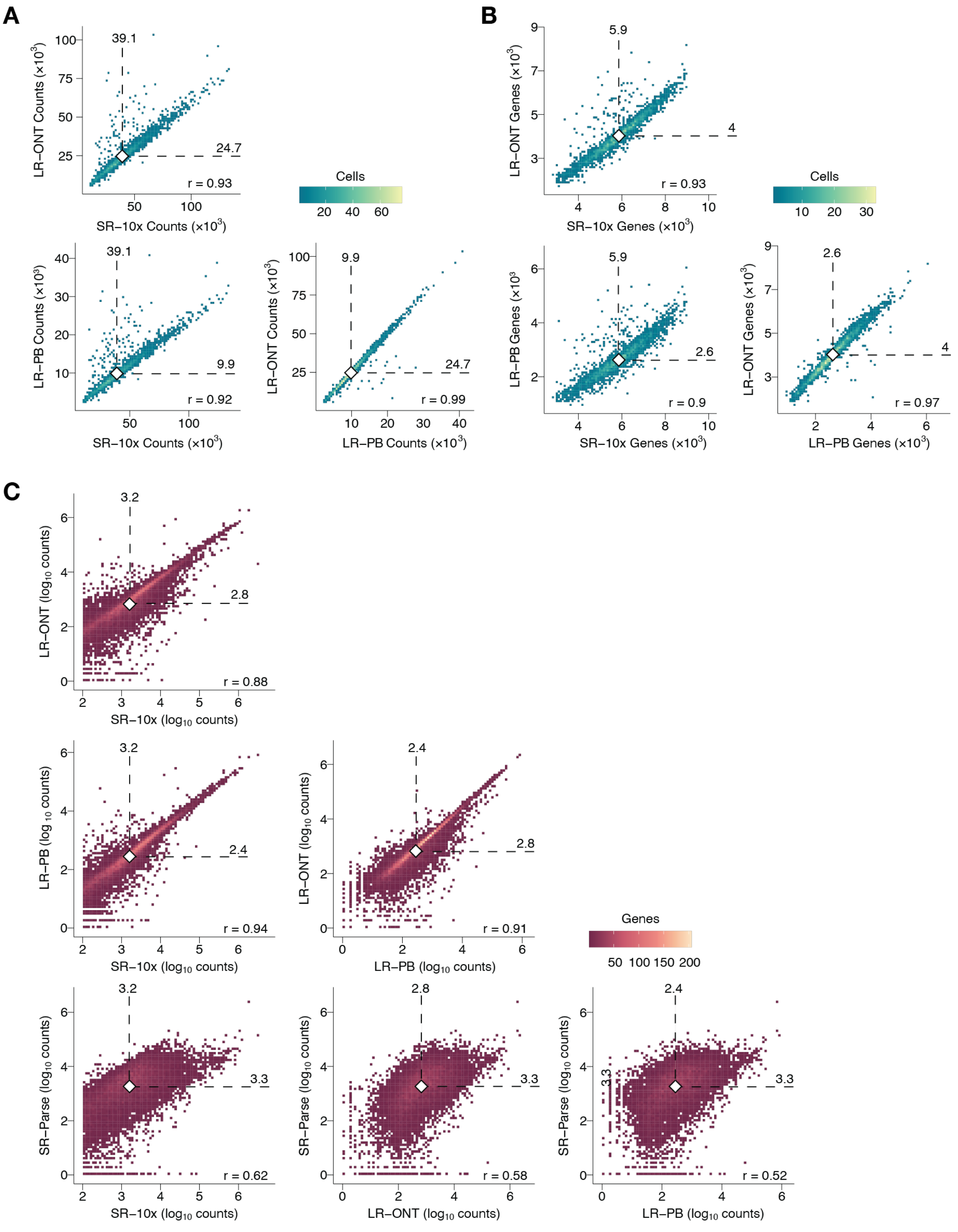


**Supplementary Figure 3. Cross-platform concordance at cell and gene level.**

**A–B.** Per-cell UMI counts (A) and detected genes (B) compared pairwise across the three 10x-based datasets, restricted to cells with matching barcodes. Color indicates local cell density. Diamond markers indicate per-axis medians; dashed lines show median values. Pearson's r shown. **C.** Gene-level expression (log10 total counts) compared pairwise across all four datasets, restricted to consensus genes. Color indicates local gene density. Diamond markers and dashed lines indicate per-axis medians. Pearson's r shown.

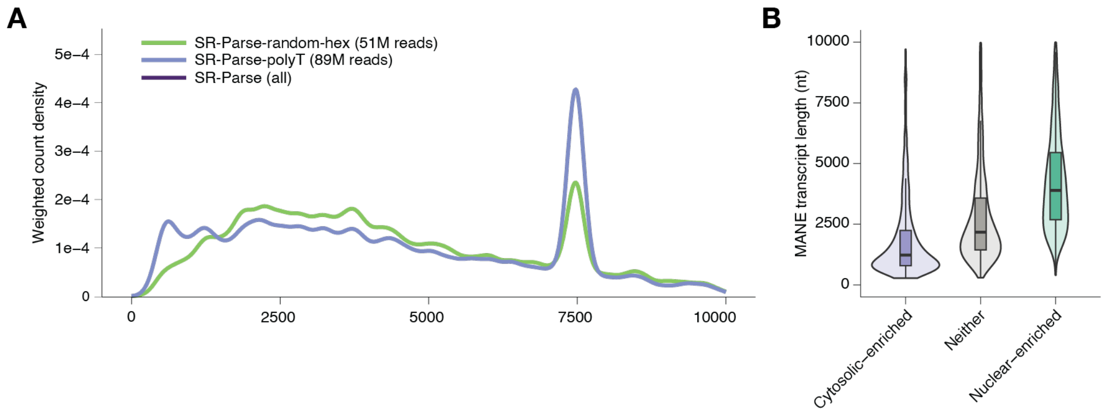


**Supplementary Figure 4. Parse priming biases and nuclear transcript enrichment.**
**A.** Normalized gene body coverage plotted across housekeeping genes for SR-Parse reads originating from random hexamer primers, oligo-dT primers, or all reads combined. Read counts per priming strategy are shown in the legend. **B.** Genes were classified as cytosolic-enriched, nuclear-enriched, or neither based on a $\log_2$ ratio of mean nuclear to cytosolic expression in K562 cells, using publicly available ENCODE cell fractionation data. Thresholds of $\log_2$ ratio $< -1$ and $> 1$ were applied for cytosolic- and nuclear-enriched genes respectively. MANE Select transcript lengths are shown for each group. All pairwise comparisons were significantly different (Mann-Whitney U test, adjusted $p < 2\times10^{-16}$).

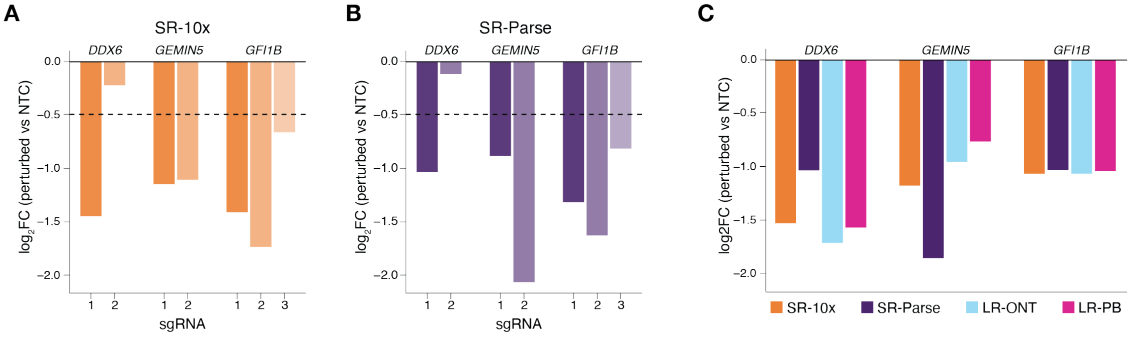


**Supplementary Figure 5. Target gene knockdowns per guide RNA.**
**A-B.** $\text{Log}_2\text{FC}$ of target gene expression (vs NTC) per sgRNA for SR-10x (**A**) and SR-Parse (**B**). Dashed line: knockdown threshold ($\log_2\text{FC} < -0.5$). **C.** $\text{Log}_2\text{FC}$ per target gene across datasets, excluding sgRNAs failing knockdown threshold.

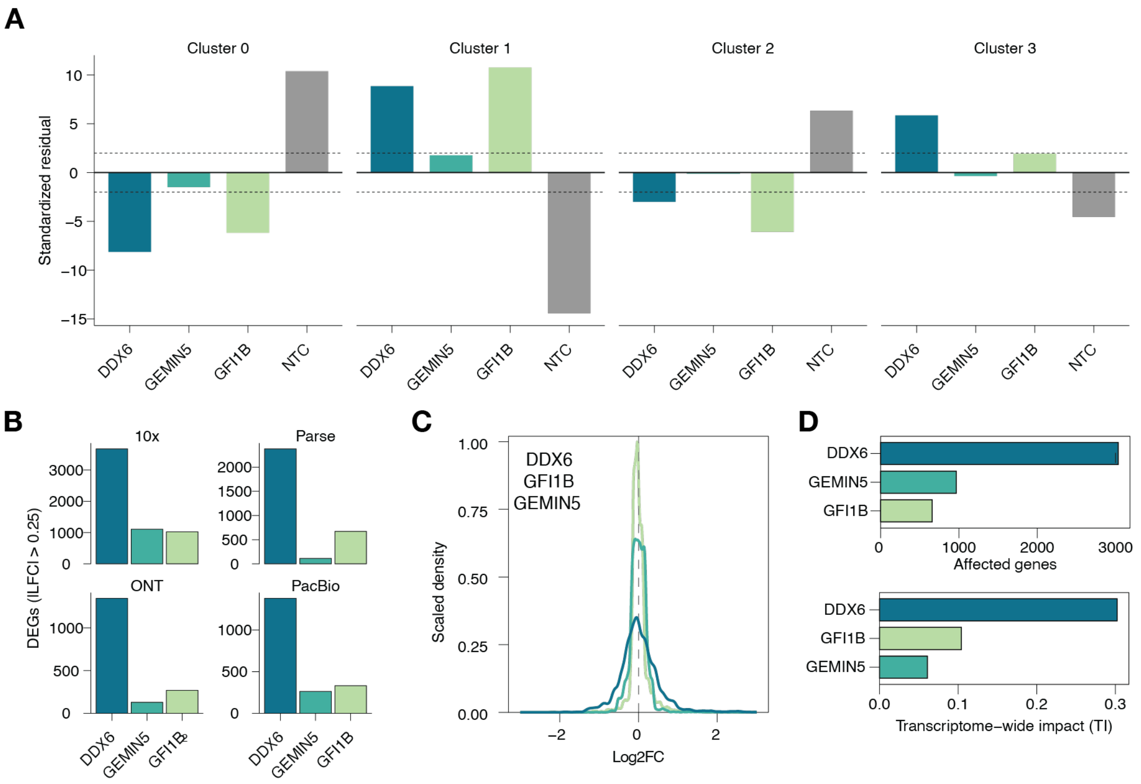


**Supplementary Figure 6. Perturbation effects on cell state and transcriptome-wide impact.**
**A.** Standardized residuals from a chi-squared test of perturbation enrichment across Seurat clusters. Dashed lines indicate ±2, the threshold for significant enrichment or depletion. **B.** Number of DEGs per perturbation and dataset (DESeq2, $|\log_2 FC| > 0.25$, FDR < 0.05). **C.** TRADE-inferred distribution of true $\log_2$FC effect sizes per perturbation in SR-10x, scaled to a maximum density of 1. Dashed line at $\log_2 FC = 0$. **D.** TRADE summary metrics for each perturbation: number of genes with non-zero true effects (top) and transcriptome-wide impact score (bottom).

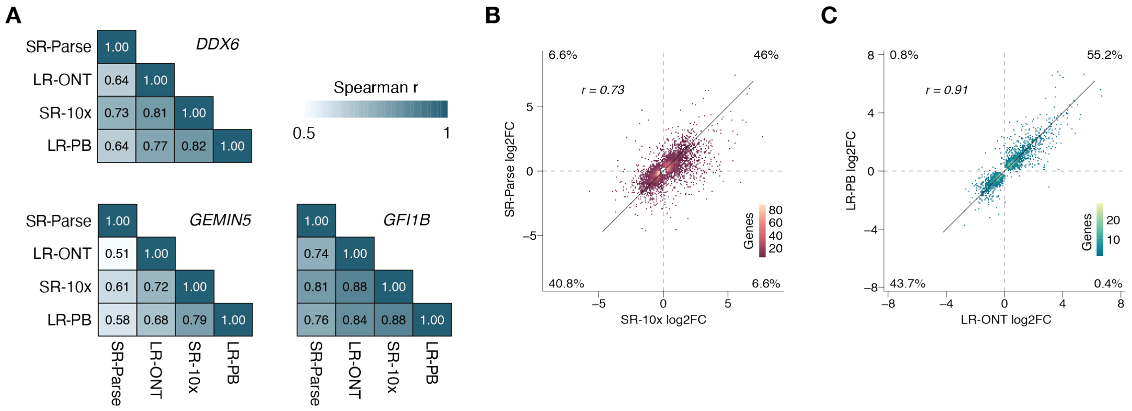


**Supplementary Figure 7. Correlation between differentially expressed genes $\log_2$FCs.**

**A.** Pairwise Spearman correlations of DEG $\log_2$FC values per perturbation across all four datasets. **B–C.** $\log_2$FC comparison between SR-10x and SR-Parse (B) and between LR-ONT and LR-PB (C), combining all perturbations. Genes included if a DEG in either dataset for a given perturbation. Color indicates local gene density. Quadrant percentages indicate the proportion of genes. Spearman ρ shown.

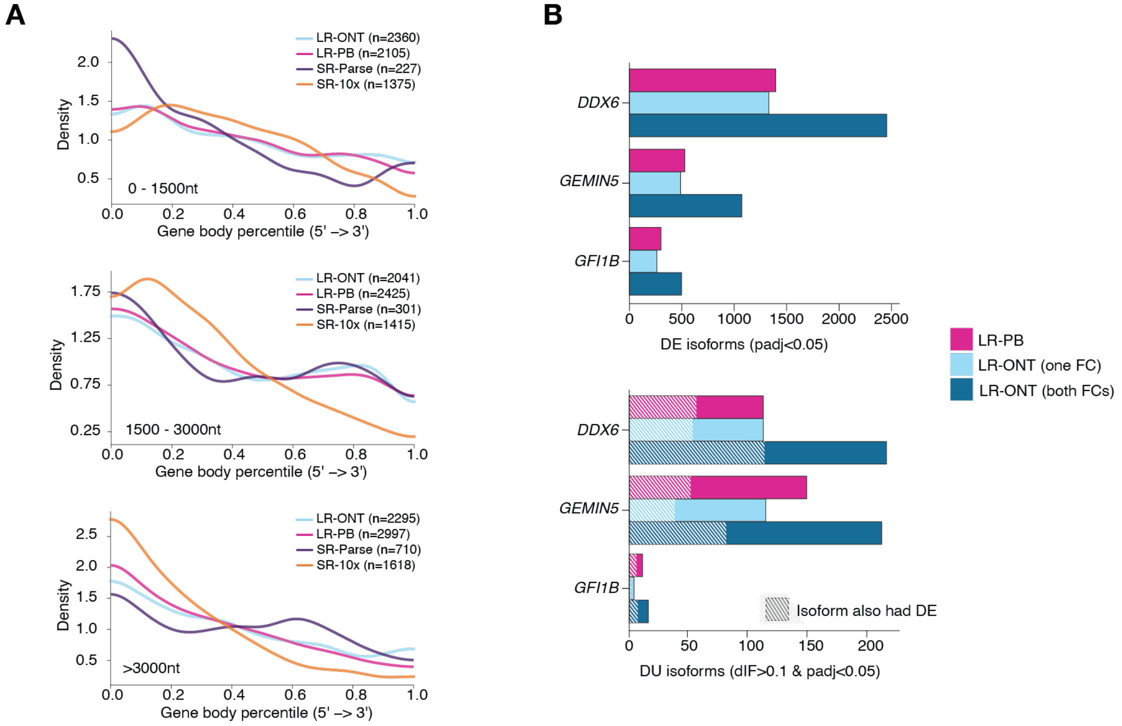


**Supplementary Figure 8. Number of differentially expressed isoforms and isoforms with differential usage.**

**A.** Density of testable splice junction positions across gene bodies, stratified by MANE Select transcript length (0–1500 nt, 1500–3000 nt, >3000 nt). The number of testable splice junctions per dataset is shown in the legend. **B.** Top: number of differentially expressed (DE) isoforms from FLAIR per perturbation and LR dataset (FDR < 0.05). Bottom: number of isoforms with significant differential usage (DU) per perturbation and LR dataset (ΔIF > 0.1, FDR < 0.05). Results are shown for LR-PB, LR-ONT with one flow cell, and LR-ONT with both flow cells combined. Hatched bars indicate DU isoforms that were also DE.

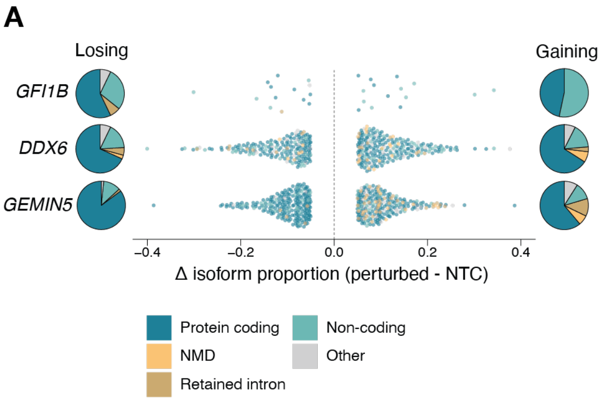


**Supplementary Figure 9. Biotypes of isoform switches.**

**A.** For each gene, the isoform with the largest decrease in usage (losing isoform) and the isoform with the largest increase in usage (gaining isoform) were selected to represent the dominant switch. The change in isoform proportion between perturbed and non-targeting control groups is shown for each perturbed gene. Colors indicate isoform biotype.

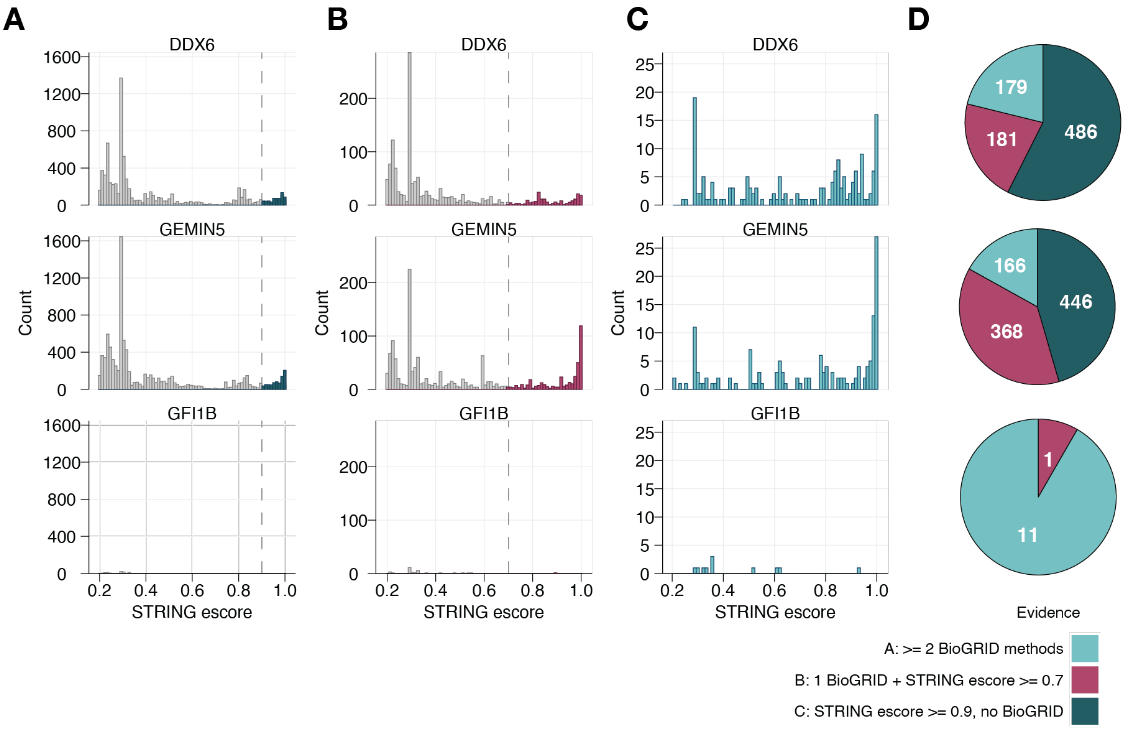


**Supplementary Figure 10. Interaction partner selection strategy using BioGRID and STRING evidence tiers.**

**A.** Distribution of STRING experimental scores for interactions with no BioGRID support. The dashed line indicates the escore threshold of 0.9 included as Tier C interaction candidates. Interactions above this threshold are highlighted. **B.** Distribution of STRING experimental scores for interactions supported by exactly one reliable BioGRID experimental method. The dashed line indicates the escore threshold of 0.7

used to define Tier B candidates. **C.** Distribution of STRING experimental scores for interactions supported by two or more reliable BioGRID experimental methods, which are retained as Tier A interactions regardless of STRING score. **D.** Composition of the final interaction set per perturbation target, showing the number of interactions contributed by each evidence tier. All panels show data for *DDX6*, *GEMIN5*, and *GFI1B* knockdown (rows).

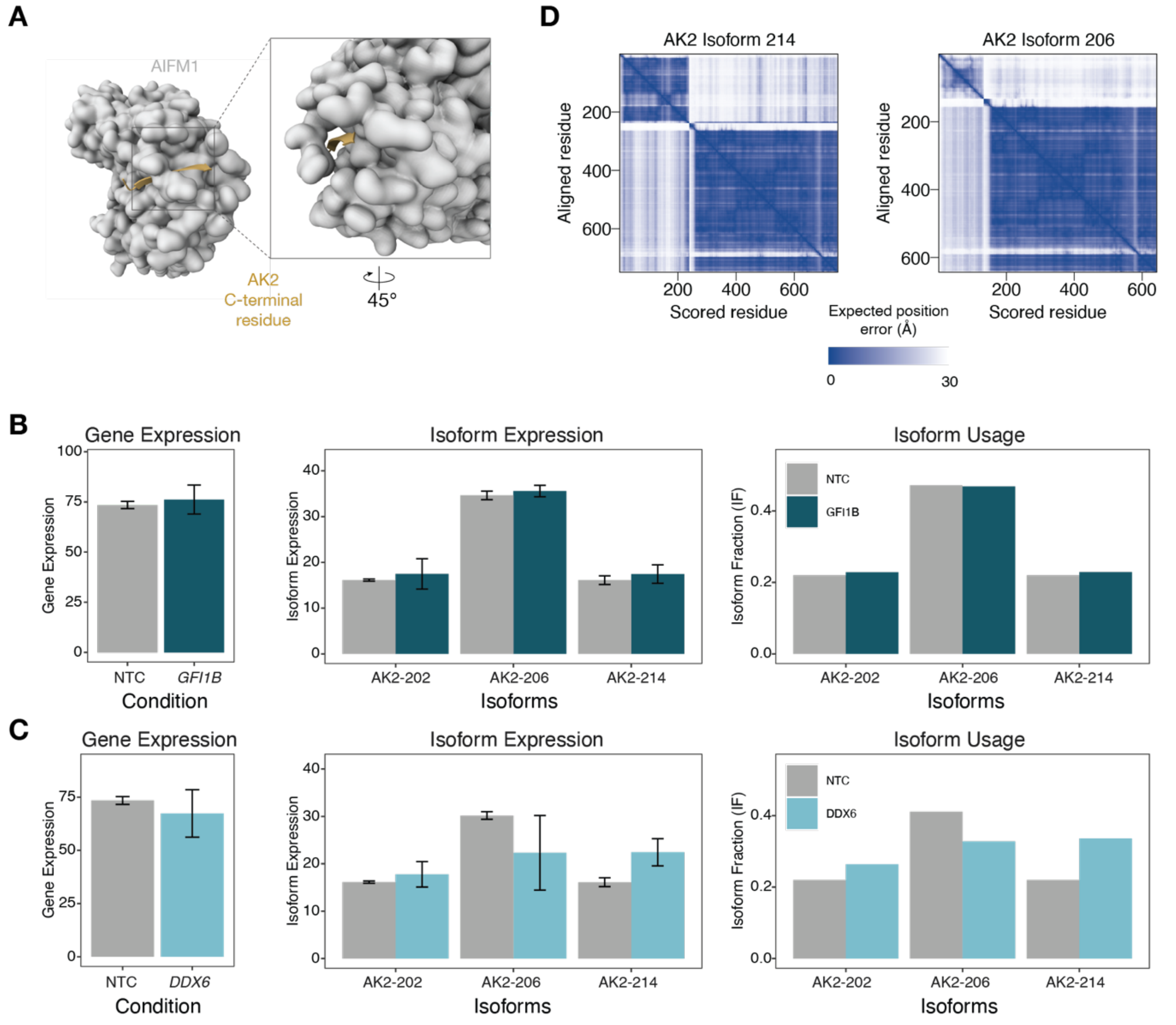


**Supplementary Figure 11. AK2 expression and isoform usage across perturbations and AK2/AIFM1 structural modeling.**

**A.** AlphaFold3-predicted structure of AIFM1 (surface) with the AK2 C-terminal residue highlighted (yellow). Inset: 45° rotation of the binding interface. **B.** AK2 gene expression (left), isoform expression (middle), and isoform fraction (right) for NTC and *GFI1B* knockdown cells. No significant changes were observed. **C.** AK2 gene expression (left), isoform expression (middle), and isoform fraction (right) for NTC and *DDX6* knockdown cells. No significant changes were observed. **D.** Predicted aligned error (PAE) plots from AlphaFold3 for AK2-214/AIFM1 (left) and AK2-206/AIFM1 (right) complexes. Lower values indicate higher confidence in relative residue positions.

**Supplementary Table 1. Cell barcodes per pseudoreplicate pseudobulk group for 10x and Parse datasets.**
**Supplementary Table 2. Gene signatures used for cell state annotation.**
**Supplementary Table 3. Differentially expressed genes across datasets from DESeq2.**
**Supplementary Table 4. Detected alternative splicing events identified with LeafCutter across datasets.**
**Supplementary Table 5. Differential isoform usage results from FLAIR for LR-ONT.**
**Supplementary Table 6. Guide RNAs used for CRISPRi of three genes and non-targeting controls.**
